# Elucidating the basis for permissivity of the MT-4 T-cell line to replication of an HIV-1 mutant lacking the gp41 cytoplasmic tail

**DOI:** 10.1101/2020.07.06.190801

**Authors:** Melissa V. Fernandez, Huxley K. Hoffman, Nairi Pezeshkian, Philip R. Tedbury, Schuyler B. van Engelenburg, Eric O. Freed

## Abstract

HIV-1 encodes an envelope glycoprotein (Env) that contains a long cytoplasmic tail (CT) harboring trafficking motifs implicated in Env incorporation into virus particles and viral transmission. In most physiologically relevant cell types, the gp41 CT is required for HIV-1 replication, but in the MT-4 T-cell line the gp41 CT is not required for a spreading infection. To help elucidate the role of the gp41 CT in HIV-1 transmission, in this study we investigated the viral and cellular factors that contribute to the permissivity of MT-4 to gp41 CT truncation. We found that the kinetics of HIV-1 production are faster in MT-4 than in the other T-cell lines tested, but MT-4 express equivalent amounts of HIV-1 proteins on a per-cell basis relative to cells not permissive to CT truncation. MT-4 express higher levels of plasma-membrane-associated Env than non-permissive cells and Env internalization from the plasma membrane is slower compared to another T-cell line, SupT1. Paradoxically, despite the high levels of Env on the surface of MT-4, two-fold less Env is incorporated into virus particles in MT-4 compared to SupT1. Cell-to-cell transmission between co-cultured 293T and MT-4 is higher than in co-cultures of 293T with most other T-cell lines tested, indicating that MT-4 are highly susceptible to this mode of infection. These data help to clarify the long-standing question of how MT-4 cells overcome the requirement for the HIV-1 gp41 CT and support a role for gp41 CT-dependent trafficking in Env incorporation and cell-to-cell transmission in physiologically relevant cell lines.

**Importance:** The HIV-1 Env cytoplasmic tail (CT) is required for efficient Env incorporation into nascent particles and viral transmission in primary CD4^+^ T cells. The MT-4 T-cell line has been reported to support multiple rounds of infection of HIV-1 encoding a gp41 CT truncation. Uncovering the underlying mechanism of MT-4 T-cell line permissivity to gp41 CT truncation would provide key insights into the role of the gp41 CT in HIV-1 transmission. This study reveals that multiple factors contribute to the unique ability of a gp41 CT truncation mutant to spread in cultures of MT-4 cells. The lack of a requirement for the gp41 CT in MT-4 is associated with the combined effects of rapid HIV-1 protein production, high levels of cell-surface Env expression, and increased susceptibility to cell-to-cell transmission compared to non-permissive cells.

## Introduction

HIV-1 Env is initially synthesized in the endoplasmic reticulum (ER) as a polyprotein precursor, gp160, which is cleaved during trafficking through the Golgi apparatus to generate the mature surface Env subunit gp120 and the transmembrane subunit gp41 (1, 2). The two subunits are non-covalently linked to form a trimeric gp120:gp41 heterodimer in the functional Env glycoprotein complex. The mature, trimeric Env complex traffics via the secretory pathway to the plasma membrane (PM), the site of viral assembly and budding. Env is expressed on the surface of infected cells and is incorporated into virus particles where it is embedded in the viral envelope.

The two subunits of Env are responsible for different functions of the glycoprotein complex. The gp120 subunit promotes particle attachment and entry by binding to receptor (CD4) and co-receptor (CXCR4 or CCR5). The gp41 subunit comprises three domains: an ectodomain that associates with gp120 and contains the determinants critical for membrane fusion, a transmembrane domain that anchors Env in the lipid bilayer, and a cytoplasmic tail (CT) that regulates a number of aspects of Env function. While the principal functions of the Env complex are well characterized, and it is clear that the gp41 CT regulates Env incorporation into virions, the precise role of the CT in Env biology remains poorly understood.

HIV-1 is transmitted to target cells *in vitro* and *in vivo* via either cell-free or cell-to-cell (C-C) infection (3, 4). Cell-free infection occurs when virions that are not associated with the virus-producing cell bind and enter uninfected target cells. C-C infection is defined as direct transmission of nascent particles at points of contact, known as virological synapses (VSs), between infected and uninfected cells. Studies have established that, *in vitro*, viral dissemination by C-C transmission is a highly efficient mode of viral transfer relative to cell-free infection (5–9). However, the relative contribution of cell-free vs C-C transmission to viral spread *in vivo* is less clear. In most cell types, viral transmission requires CT-dependent localization of Env to viral assembly sites (10–13) and Env binding to CD4 and co-receptor. A hallmark of C-C spread is the accumulation of viral proteins, in particular Gag and Env, at the VS (5, 6, 11, 14). How Env is directed to the VS is not well understood; further elucidation of this process is fundamental to our ability to design therapies capable of blocking C-C transmission.

The lentiviral gp41 CT is very long compared to those of other retroviruses; it contains 150 amino acids in the case of HIV-1 and harbors trafficking motifs implicated in Env recycling, incorporation, and viral transmission. The gp41 CT contains a highly conserved YxxΦ motif (with Φ representing a hydrophobic amino acid) known to interact with host cell clathrin adaptor protein complex 2 (AP-2) and mediate fast internalization of Env via clathrin-mediated endocytosis (2). The gp41 CT contains several other well-conserved tyrosine and dileucine motifs that may also play a role in Env trafficking and subcellular localization (2). The high degree of conservation in both the length of the gp41 CT and the YxxΦ motif suggests that these features play key roles in viral transmission. It is currently unclear whether Env recycling from the PM prior to incorporation into the assembling Gag lattice is a requisite step in Env incorporation.

Recent evidence suggests a role for recycling in Env incorporation (15–17) and many studies have explored the role of trafficking motifs in the gp41 CT in promoting the proper spatio-temporal localization of Env during assembly (1, 2, 18, 19), but the role of Env recycling in Env incorporation is not well-defined.

Wild-type (WT) HIV-1 has an average of ~10 Env trimers per virion (20) and truncation of the gp41 CT generally results in a 10-fold decrease in Env incorporation in physiologically relevant cell types (which we refer to as being non-permissive to gp41 CT truncation) (21). The sparsity of Env on HIV-1 particles suggests that Env incorporation is tightly regulated. The degree of regulation seems to be cell-type and CT dependent. For example, in the non-permissive T-cell line CEM-A, WT Env is localized at the neck of the budding particle while truncation of the gp41 CT results in a more uniform Env distribution around the virus particle (19). In the permissive COS7 fibroblast-like cell line, both WT and CT-truncated Env are evenly distributed around the virus particle. CT-dependent endocytosis of WT Env from the PM is more active in CEM-A than in COS7, suggesting that Env recycling regulates Env distribution on the virus particle. Importantly, in both cell lines, incorporation of the gp41 CT-truncated mutant was reduced compared to the WT, consistent with the essential role for the CT in trapping Env in the assembling Gag lattice (22). These data highlight the multifaceted role of the gp41 CT in regulating Env incorporation and the cell-type dependent utilization of the CT in viral assembly.

A functional interaction between the gp41 CT and the matrix (MA) domain of the Gag polyprotein has been postulated to play an important role in capturing Env during viral assembly (23, 24). Compelling evidence for the trapping of the gp41 CT by the Gag lattice has been previously demonstrated by various studies showing the clustering of Env trimers at sites of virus assembly (10, 12, 19, 22) and retention of full-length Env, but not CT-truncated Env, in detergent-stripped Gag virus-like particles (25). Further evidence for the trapping of the gp41 CT by the Gag lattice is provided by the ability of single amino acid changes in MA to block WT Env incorporation (26–33). Truncation of the gp41 CT reverses the Env incorporation block imposed by these point mutations in MA (26, 29, 31, 33). Similarly, Env incorporation is inhibited by small intragenic deletions in MA or deletion of the globular domain of MA, and this inhibition is relieved by truncation of the gp41 CT (31, 34). Highlighting the importance of MA-gp41 CT interactions, a small deletion in the gp41 CT that inhibits Env incorporation is rescued by a single amino acid change in MA (35). Further underscoring the intimate relationship between MA and the gp41 CT, truncation of the gp41 CT abrogates Gag’s ability to repress premature fusion of Env with the target cell membrane (12, 36, 37). Recent studies have demonstrated an important role for trimerization of the MA domain of Gag in the formation of a Gag lattice that accommodates the long gp41 CT during Env incorporation (27, 28, 38). While evidence of the relationship between the gp41 CT and MA is well appreciated, the precise mechanism of how Env is incorporated into the assembling Gag lattice is not well understood.

Our understanding of the function of the gp41 CT and MA in Env incorporation suggests four general models of Env incorporation: 1) passive incorporation (Env incorporation does not require its concentration at assembly sites), 2) Gag-Env co-targeting (Gag and Env are both targeted to an assembly platform, such as a membrane microdomain), 3) direct Gag-Env interaction (Gag directly binds the gp41 CT, thereby capturing it into assembling virions), and 4) indirect Gag-Env interaction (a host factor serves as a bridge between Gag and Env for the capture of Env by the assembling Gag lattice) (1, 23). These models are not mutually exclusive; for example, Env could colocalize with Gag at sites of virus assembly and then be captured by the Gag lattice via direct interactions between MA and the gp41 CT, and other combinations of these models can be readily envisioned. Interestingly, even foreign viral glycoproteins have been shown to cluster at HIV-1 assembly sites in the context of pseudotype particle formation (39, 40).

The unusual ability of certain T-cell lines (notably MT-4, C8166, and M8166) to be highly susceptible to HIV-1 infection and permissive to loss of certain viral protein functions is incompletely understood. In particular, MT-4 are known to be permissive, relative to other T-cell lines, to deletion of the gp41 CT (14, 21, 29, 31, 33, 38, 41–45), loss of integrase (IN) function (46, 47), disruption of proper capsid (CA) multimerization (48), deletion of Nef (49), and a large MA deletion (34). A functional IN protein is required for productive HIV-1 replication in a majority of cell lines and human peripheral blood mononuclear cells (hPBMCs) (47). MT-4 and C8166 are known to be permissive to type I IN mutations (mutations in IN that specifically block viral DNA integration into the host cell genome) while Jurkat E6.1 and hPBMCs are non-permissive to defects in integration (47). It was recently reported that MT-4 and C8166 are likely permissive to type I IN mutations due to HTLV-I Tax expression inducing NF-κB protein recruitment to the HIV-1 LTR on unintegrated HIV-1 DNA (46). Furthermore, it was previously suggested by Emerson et al. (14) that HTLV-I Tax expression may contribute to MT-4 permissivity by causing potent and chronic activation of NF-κB signaling (50), resulting in transactivation of the HIV-1 LTR (51). The ability of MT-4 to support rapid replication of HIV-1 and gain second-site compensatory mutations unlikely to be acquired in less-permissive cells has led to the use of this cell line for selection experiments with a wide variety of defective MA and CA mutants (29, 48, 52–54). Uncovering the underlying mechanisms of MT-4 cell line permissivity is important because of the frequent use of this cell line in HIV-1 replication studies (13, 55–57).

In this study, we examined a number of factors involved in viral replication and spread in cells both permissive (i.e., MT-4) and non-permissive (i.e., all other T-cell lines tested) for gp41 CT truncation. Our results show that HTLV-I Tax expression is not the sole determinant of permissivity to gp41 CT truncation. Rather, high surface Env expression, rapid kinetics of HIV protein production, and efficient C-C transmission likely contribute to the MT-4 permissivity to CT-truncation.

## Results

### Lineage and validation of cell lines used to interrogate T-cell line permissivity to replication of an HIV-1 mutant lacking the gp41 CT

To investigate the mechanistic basis for the permissivity of MT-4 cells to replication of a gp41 CT-truncation mutant, a panel of five T-cell lines were selected based on their origins (Fig. 1) and previously reported permissivity, or lack thereof, to gp41 CT truncation. Previous studies have reported that MT-4 (14, 21, 29, 31, 33, 38, 41–45) and M8166 (58) are permissive to truncation of the gp41 CT, and it is established that hPBMCs, SupT1, and Jurkat E6.1 are not permissive (14, 21). Therefore, two HTLV-1^-^ lymphoma-derived cell lines, SupT1 (59) and Jurkat E6.1 (60) (Fig. 1A-B); and three HTLV-1-transformed lines, MT-4 (61), C8166 (62), and M8166 (Fig. 1C-D) were selected for this study. The cell line lineage and HTLV-I particle and RNA production status of each cell line are shown in Fig 1. C8166, M8166, and MT-4 all express HTLV-I RNA but do not produce viral particles. Due to the presence of HTLV-I RNA, they do express the HTLV-I Tax protein (43, 63, 64). SupT1 and Jurkat E6.1 do not express HTLV-I RNA or proteins (59, 60).

**Figure 1.**
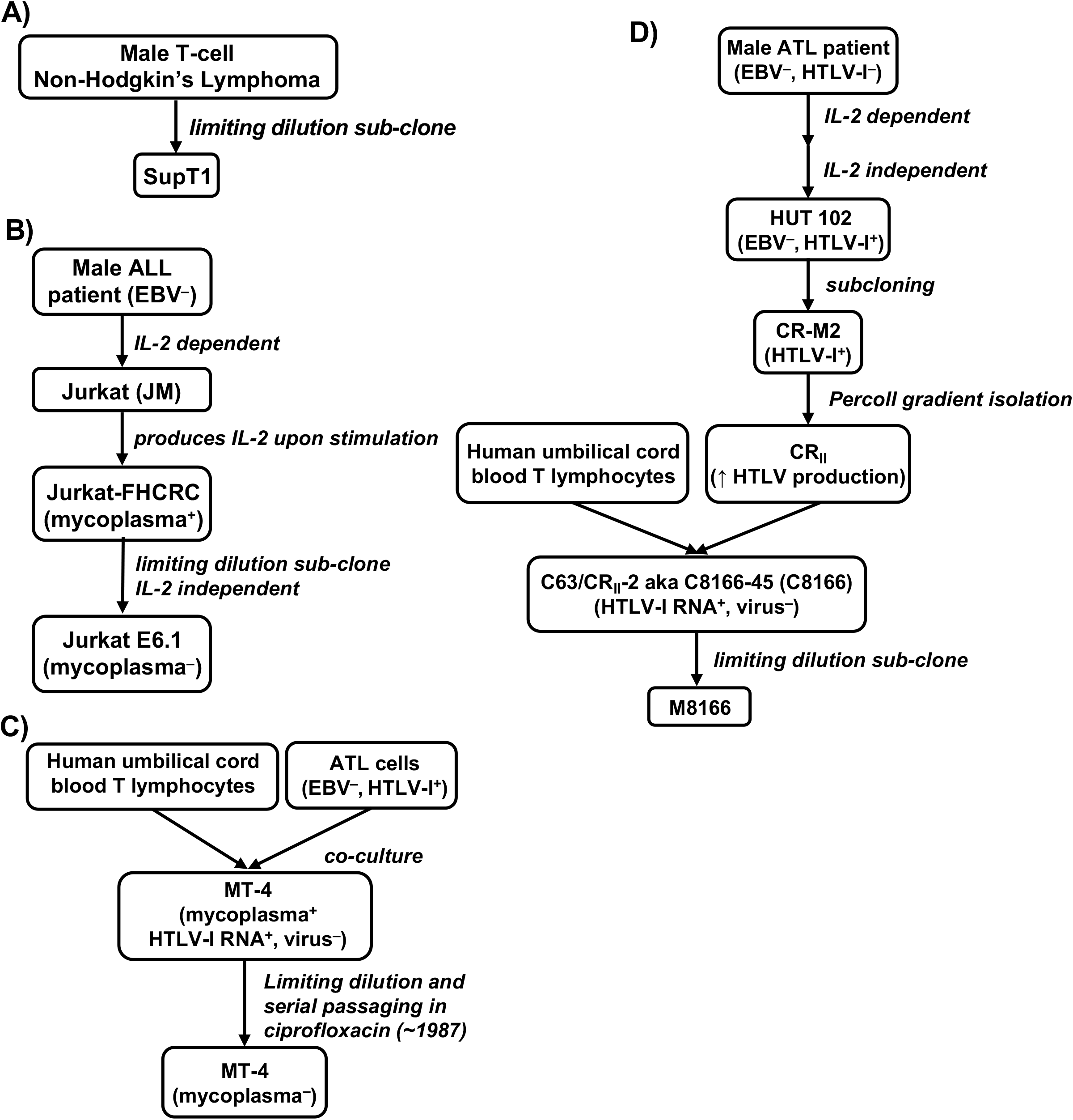
Lineage of cell lines used to interrogate T-cell line permissivity to replication of an HIV-1 mutant lacking the gp41 CT. A) The SupT1 T-cell line was derived from a pleural effusion of a male patient with non-Hodgkin’s lymphoma. Cells were sub-cloned by limiting dilution to generate a cell line capable of continual growth from a single-cell colony (59). B) The peripheral blood of a 14-year-old boy with relapsed acute lymphocytic leukemia (ALL) was used to generate the EBV^-^, IL-2-dependent JM T-cell line (60). The JM line was then subcloned to generate the Jurkat-FHCRC subclone for its ability to produce IL-2 upon stimulation with phorbol esters or lectins (99). “-FHCRC” designation indicates the cell line originated at the Fred Hutchinson Cancer Research Center. Jurkat-FHCRC was then subjected to limiting dilution to generate an IL-2-independent, mycoplasma-free cell line, Jurkat E6.1 (100). C) To generate the MT-4 cell line, cells from an adult male ATL patient were co-cultured with male human infant cord leukocytes, therefore it is unknown whether these cells are of cord leukocyte or ATL cell origin (61). Cells were gifted to the lab of Douglas Richman by Harada et al. (101) and serially passaged by terminal dilution cloning of the cells in the presence of ciprofloxacin until they were determined to be mycoplasma free. These cells were then donated to the NIH ARP by Douglas Richman (D. Richman, personal communication). D) To generate the C8166 T-cell line, cells were first acquired from a 26-year-old male ATL patient’s inguinal lymph node (102) and maintained in IL-2 for several passages until they were deemed IL-2 independent (103). This T-cell line, HUT 102, was determined to be EBV^-^ and HTLV^+^. The CR-M2 subclone of HUT 102 was then isolated and further purified by percoll gradient isolation to generate a cell line with increased HTLV production, designated CRII (104). CRII was then used to transform human umbilical cord blood T leukocytes by cell hybridization to generate the C63/CRII-2 cell line, also known as the C8166-45 (aka 81-66 or C8166) cell line (62). The “-45” designation indicates the cell line has 45 chromosomes. “CR” indicates the cells were transformed by the HTLV-I_CR_ virus. Characterization of this line found that it produces HTLV-I RNA but no virus particles (62). C8166 were then subjected to limiting dilution to generate a clone more susceptible to formation of syncytia when cultures were infected with HIV-1 (105). This new C8166-derived cell line was named M8166.

Short tandem repeat (STR) profiling was utilized to validate the identity of all the cell lines in our T-cell panel (Table 1). The MT-4 STR profile, recently validated by functional assays, morphological analysis, and assessment of HTLV-I Tax protein expression (43), was an exact match to the Cellosaurus reference profile (Table 1). Jurkat E6.1 and SupT1 STR profiles were both close matches to the published STR profiles. The STR profile revealed some genetic instability in the SupT1 line compared to SupT1-CCR5, indicated by the presence of extra, lower-intensity alleles at several gene loci (data not shown). Overall, the STR profile of our SupT1 cell line was a 95% match to the SupT1-CCR5 line, confirming the identity of our laboratory SupT1 line. The Jurkat E6.1 line also displayed some genetic instability (data not shown), which accounts for its imperfect match to the Cellosaurus reference profile.

**TABLE 1.** T-cell Line STR profiles

| **Genetic Loci** | **Jurkat E6.1** | **MT-4** | **C8166 & M8166** | **SupT1** |
| --- | --- | --- | --- | --- |
| D3S1358 | 15, 17 | 17 | 15,16 | 16, 17, 18, 19 |
| TH01 | 6, 9.3 | 7 | 6, 9.3 | 9.3 |
| D21S11 | 30.2, 31.2, 33.2 | 28, 30.3 | 27, 30 | 28, 31, 32, 33 |
| D18S51 | 12, 13, 20, 21 | 13 | 14, 17 | 13, 14 |
| Penta E | 10, 12 | 5, 15 | 7, 11 | 13, 14, 16 |
| D5S818 | 9 | 10, 11 | 12 | 11 |
| D13S317 | 8, 12 | 12 | 10, 11 | 10, 11, 12 |
| D7S820 | 8, 12 | 8, 10 | 9, 10 | 11 |
| D16S539 | 11 | 9, 12 | 12, 13 | 9, 12 |
| CSF1PO | 10, 11, 12, 13 | 11, 12 | 10 | 10, 11, 12 |
| Penta D | 11, 13 | 10, 13 | 11, 15 | 12 |
| vWA | 18, 19 | 17, 18 | 15, 16 | 16,17,18,19 |
| D8S1179 | 12, 13, 14, 15 | 10, 15 | 11, 14 | 13,14 |
| TPOX | 8, 10 | 11 | 11 | 9 |
| FGA | 20, 21 | 23 | 21, 22 | 19, 20, 21 |
| AMEL | X,Y | X,Y | X,Y | X,Y |
| % Match to<br>Cellosaurus<br>reference | 87.88 | 100 | 98.31 | 82.35 |
| source | NIH-ARP | NIH-ARP | NIH-ARP | unknown |
| catalog no. | 177 | 120 | 404 & 11395 |  |

M8166 is a subclone of C8166 (Fig. 1D), and therefore these two lines share the same STR profile (Table 1), which was a ~98% match to the cell line profile published in the Cellosaurus database and in a separate report (65). To further confirm the identity of the M8166 and C8166 cell lines, they were assessed for their capacity to host replication of WT SIVmac239 (Fig. 2). The C8166 line has been reported to exhibit a block to efficient SIV replication (66), whereas M8166 is reportedly capable of hosting multiple rounds of SIVmac239 replication (67). Consistent with these reports, SIVmac239 did not establish a spreading infection in C8166 (Fig. 2a) while it did in M8166 (Fig. 2B). Together, the STR profile and capacity of M8166, but not C8166, to support multiple rounds of SIVmac239 replication confirm the identity of the C8166 and M8166 cell lines used in this study.

**Figure 2.**
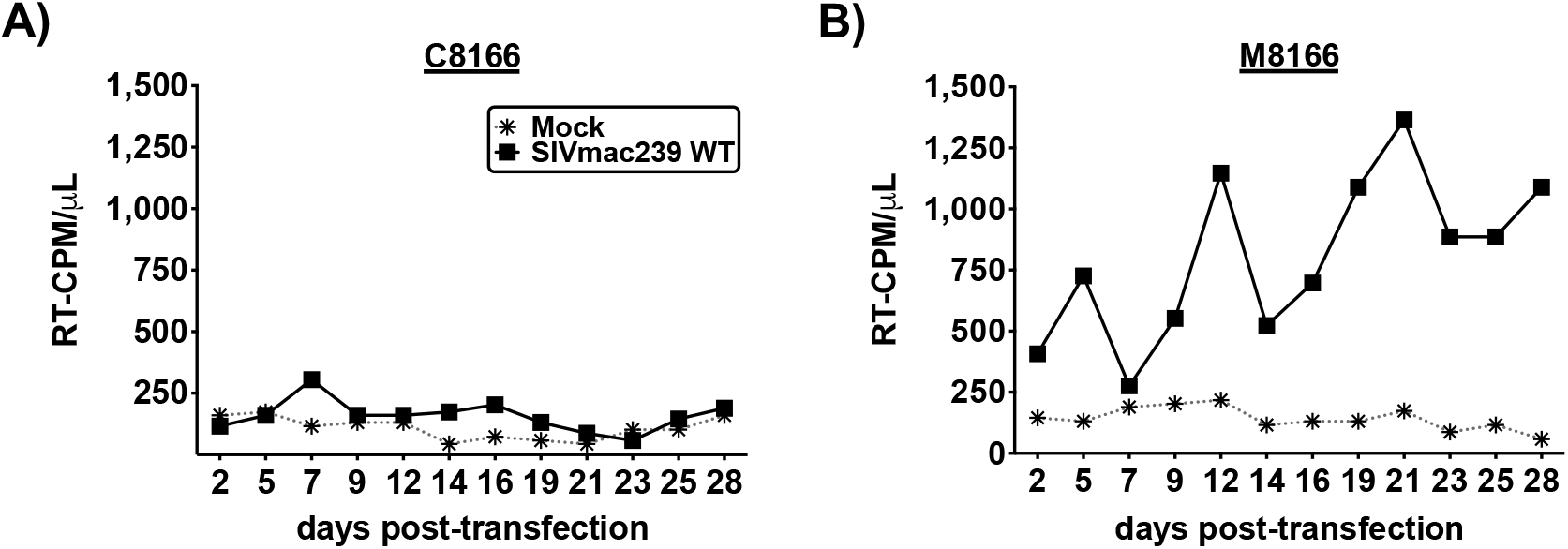
Spreading infection kinetics of SIVmac239 in C8166 and M8166. A) C8166 and B) M8166 were transfected with SIVmac239 encoding the indicated Env CT genotype. Supernatant was sampled every 2-3 days for analysis by HIV-1 RT assay and cell cultures were split 1/2. Data are representative of 3 independent experiments.

### MT-4 is the only T-cell line tested in this study that is permissive to the gp41 CT truncation mutant CTdel144

HIV-1 replication in most T-cell lines is abrogated by truncation of the gp41 CT (14, 21), but several studies have demonstrated that the MT-4 cell line is able to propagate a gp41 CT-deleted mutant (14, 21, 29, 31, 33, 38, 41–45). To evaluate the ability of the CTdel144 mutant, which lacks 144 amino acids from the gp41 CT (26), to replicate in HTLV-I-transformed T-cell lines, MT-4, C8166 and M8166 cells were transfected with the WT pNL4-3 molecular clone or the CTdel144 derivative, to initiate a spreading infection (Fig. 3A). MT-4, C8166, and M8166 (Fig. 3B-D) supported rapid WT HIV-1 replication with supernatant HIV-1 reverse transcriptase (RT) activity detectable within 3-5 days post-transfection (Fig. 3B). Replication of HIV-1 in T-cell lines not permissive to truncation of the gp41 CT - or in phytohemagglutinin (PHA)-activated PBMCs isolated from healthy hPBMCs - was markedly slower than in MT-4 cells (Fig. 3E-G compared to Fig. 3B). Consistent with previous reports (14, 21, 29, 31, 33, 38, 41–45), the MT-4 cell line also supported replication of the CTdel144 mutant. In contrast, neither of the other two HTLV-1-transformed T-cell lines (C8166 and M8166), the two HTLV-1^-^ lymphoma-derived cell lines (SupT1 and Jurkat E6.1), or hPBMCs supported replication of CTdel144 (Fig. 3B compared to Fig. 3C-G). Therefore, under these standard conditions, MT-4 is unique among the T-cell lines tested here in their capacity to support multiple rounds of CTdel144 replication.

**Figure 3.**
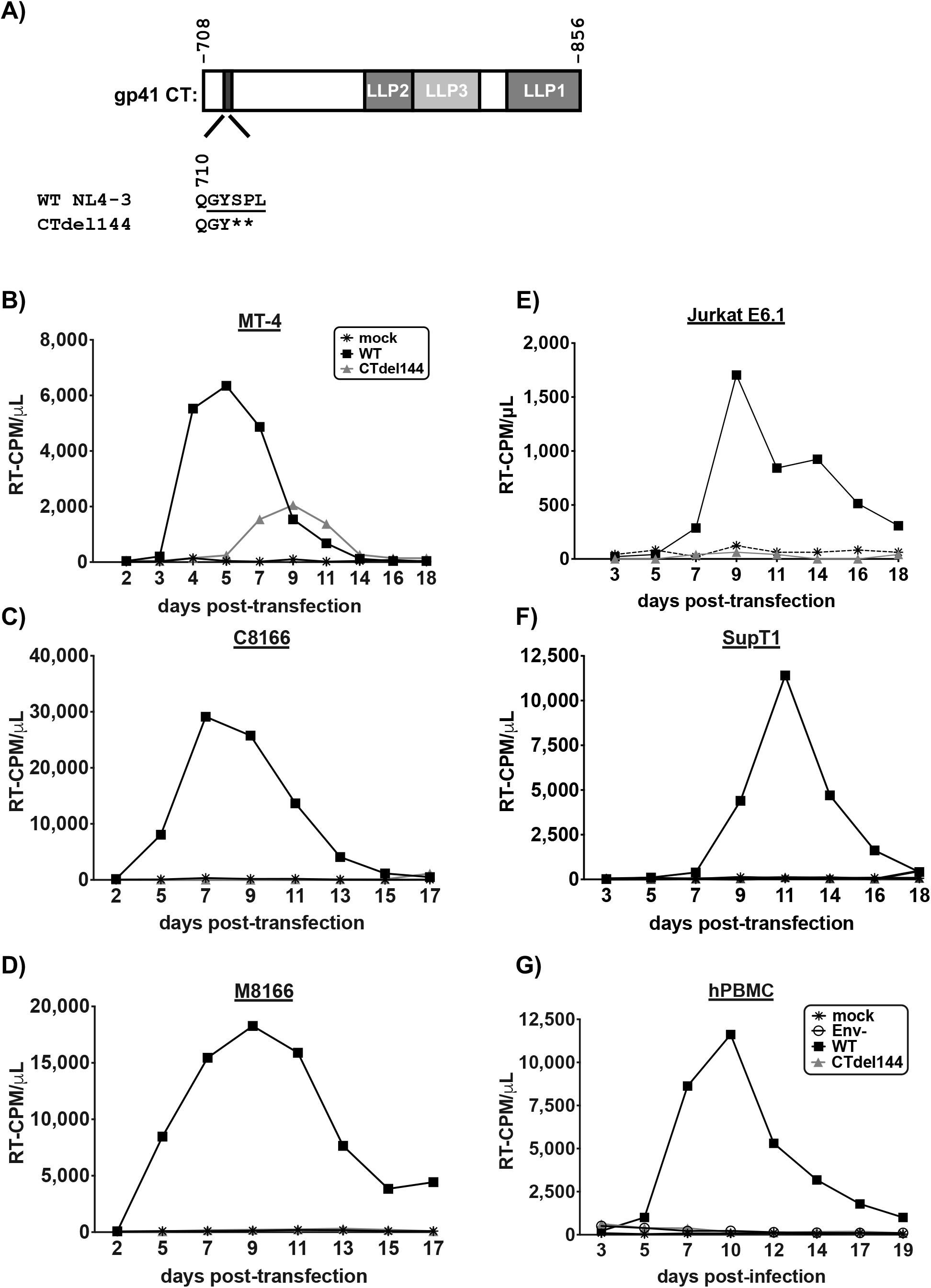
MT-4 is the only T-cell line tested that is permissive to the gp41 CT truncation mutant CTdel144. A) Schematic representation of the NL4-3 gp41 CT and gp41 CT genotypes used in this study. The lentiviral lytic peptide (LLP) domains are indicated in grey boxes and the highly conserved tyrosine endocytosis motif is indicated with a shaded black rectangle. The CTdel144 mutant was generated by introducing two stop codons in the highly conserved tyrosine endocytosis motif, resulting in a CT of 4 amino acids (26). The numbers above the gp41 CT schematic indicate the first and last amino acid positions of the gp41 CT. The number below the gp41 CT indicates the position of the QGYSPL sequence. The tyrosine endocytosis motif, GYSPL, is underlined. (B-G) Replication curves for spreading infection are shown. B-F) Cell lines were either mock transfected or transfected with pNL4-3 encoding the indicated gp41 CT genotype. Cells were split B-D) 1/2 or E-F) 1/3 every 2-3 days, and an aliquot of the supernatant was reserved for analysis of HIV-1 RT activity at each time point. G) hPBMCs were transduced with VSV-G-pseudotyped NL4-3 encoding the indicated gp41 CT genotype or mock infected. Supernatant was sampled every 2-3 days for analysis of HIV-1 RT activity and cell cultures were supplemented with fresh media. Data are representative of 3 independent experiments (B-F) and 3 donors (G).

### NF-κB-target gene expression in Tax-expressing T-cell lines is higher than in SupT1

To determine the relative levels of Tax expression in MT-4, C8166 and M8166, western blot analysis of cell lysates was performed (Fig. 4A). To control for antibody specificity, two sets of controls were included: 1) a panel of adult T-cell leukemia (ATL) cells previously reported to have lost Tax expression (68), and 2) 293T cells transfected with a Tax expression vector (Fig. 4A). Western blot analysis indicated that C8166 expressed substantially more Tax protein than MT-4 or M8166, while a small amount of Tax expression in the ATL-55T cell line was detectable.

**Figure 4.**
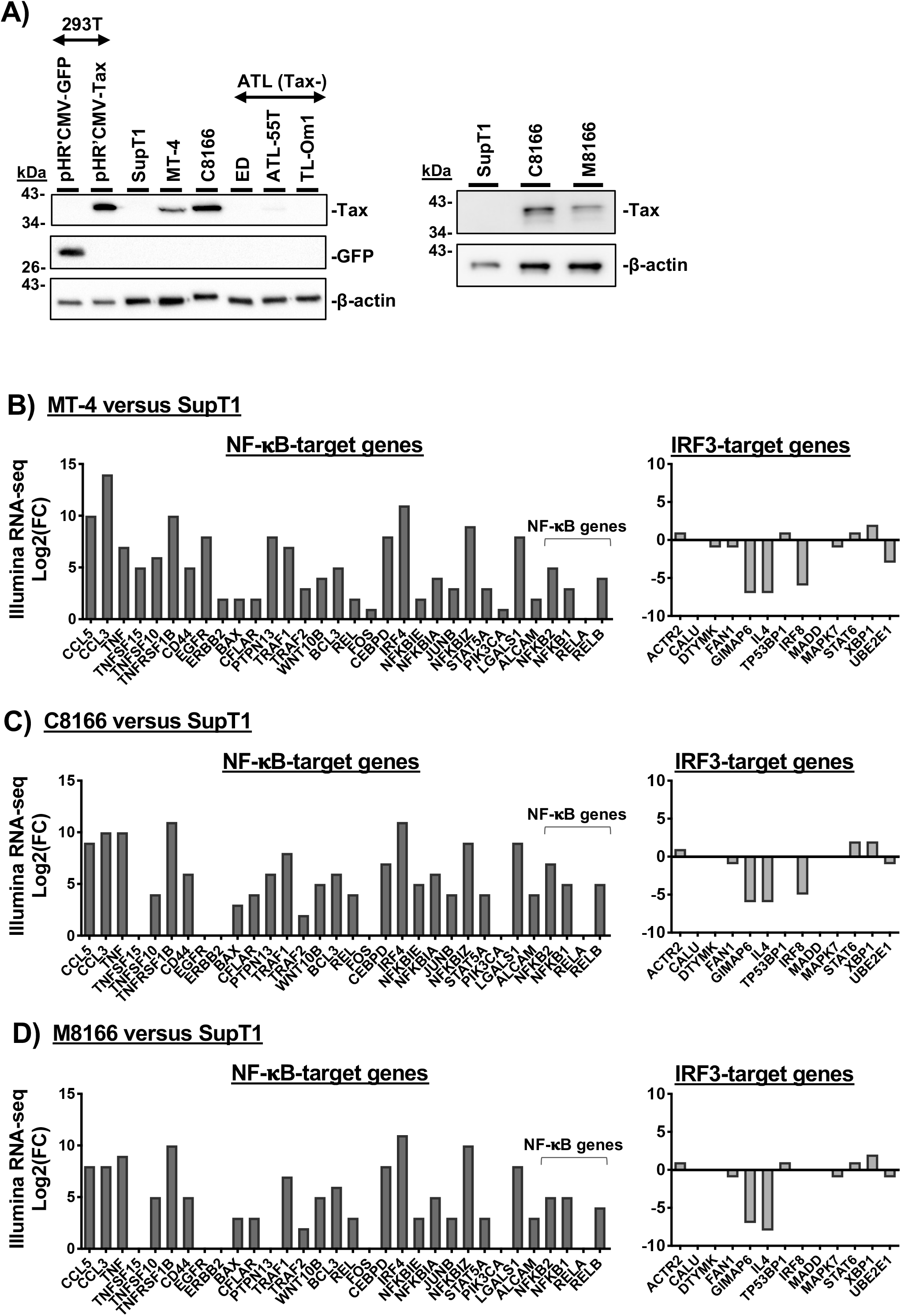
NF-κB-target gene expression in Tax-expressing T-cell lines is higher than in SupT1. A) Cell lysates from the indicated cell lines were analyzed by western blot for HTLV-I Tax expression. 293T were transfected with a GFP expression vector as a transfection control, or a Tax expression vector as a control for antibody specificity. ATL Tax-deficient cells were included as negative control for Tax expression in immortalized T-cell lines. B-D) RNA levels of the indicated NF-κB- and IRF3-target genes were compared to SupT1. RNA levels were determined by Illumina RNA-seq. As described in more detail in the methods, comparisons are reported as the Log2 of the fold-change (FC) of the HTLV-transformed line (MT-4, C8166, or M8166) relative to the lymphoma-derived T-cell line, SupT1.

Tax is a known oncogene whose expression induces rapid senescence through potent and persistent NF-κB hyperactivation and upregulation of NF-κB-regulated genes (68, 69). To ascertain whether Tax expression in these cells has resulted in hyperactivation of NF-κB signaling, Illumina RNA-seq was employed to measure the RNA expression of NF-κB-dependent and -independent genes in untreated cells. A panel of NF-κB target genes was selected from a database of genes activated by NF-κB; these included immunoreceptor genes and genes involved in proliferation, apoptosis, stress response, and cytokine-stimulation (70). A control panel of NF-κB-independent genes targeted by the IRF3 transcription factor was derived from the ENCODE Transcription Factors database (available on the Harmonizome search engine) for IRF3-target genes identified by ChIP-seq (71, 72). IRF3-target genes also targeted by NF-κB, identified by their presence in the NF-κB-target genes panel (70), were not used in the IRF3-dependent target gene analysis. Overall, NF-κB-target gene transcripts were upregulated in all three HTLV-I-transformed lines relative to SupT1 (Fig. 4B-D - left). IRF3-target genes were not consistently up- or down-regulated (Fig. 4B-D - right), consistent with a Tax-dependent chronic hyperactivation of NF-κB, but not IRF3, signaling in Tax-expressing cell lines. These data establish that Tax transactivation of the HIV-1 LTR is not solely responsible for MT-4 permissivity to gp41 CT truncation.

### Viral entry mediated by full-length and CT-truncated Env is equivalent among HTLV-transformed T-cell lines

Binding of virion-associated Env to the CD4 receptor and CXCR4 or CCR5 co-receptor is an essential step for infection of human T cells by most strains of HIV-1. T-cell line tropic isolates, like NL4-3, use CXCR4 as their coreceptor. To determine whether high-level expression of CD4 or CXCR4 on the surface of MT-4 cells could contribute to the permissivity of this cell line to CT-truncated HIV-1, surface CD4 and CXCR4 was measured on the panel of T-cell lines (Fig. 5A-B). While MT-4 cells did express more CD4 and CXCR4 than C8166 or M8166, they expressed less CD4 than SupT1. Therefore, receptor and coreceptor expression levels do not account for the permissive phenotype exhibited by MT-4.

**Figure 5.**
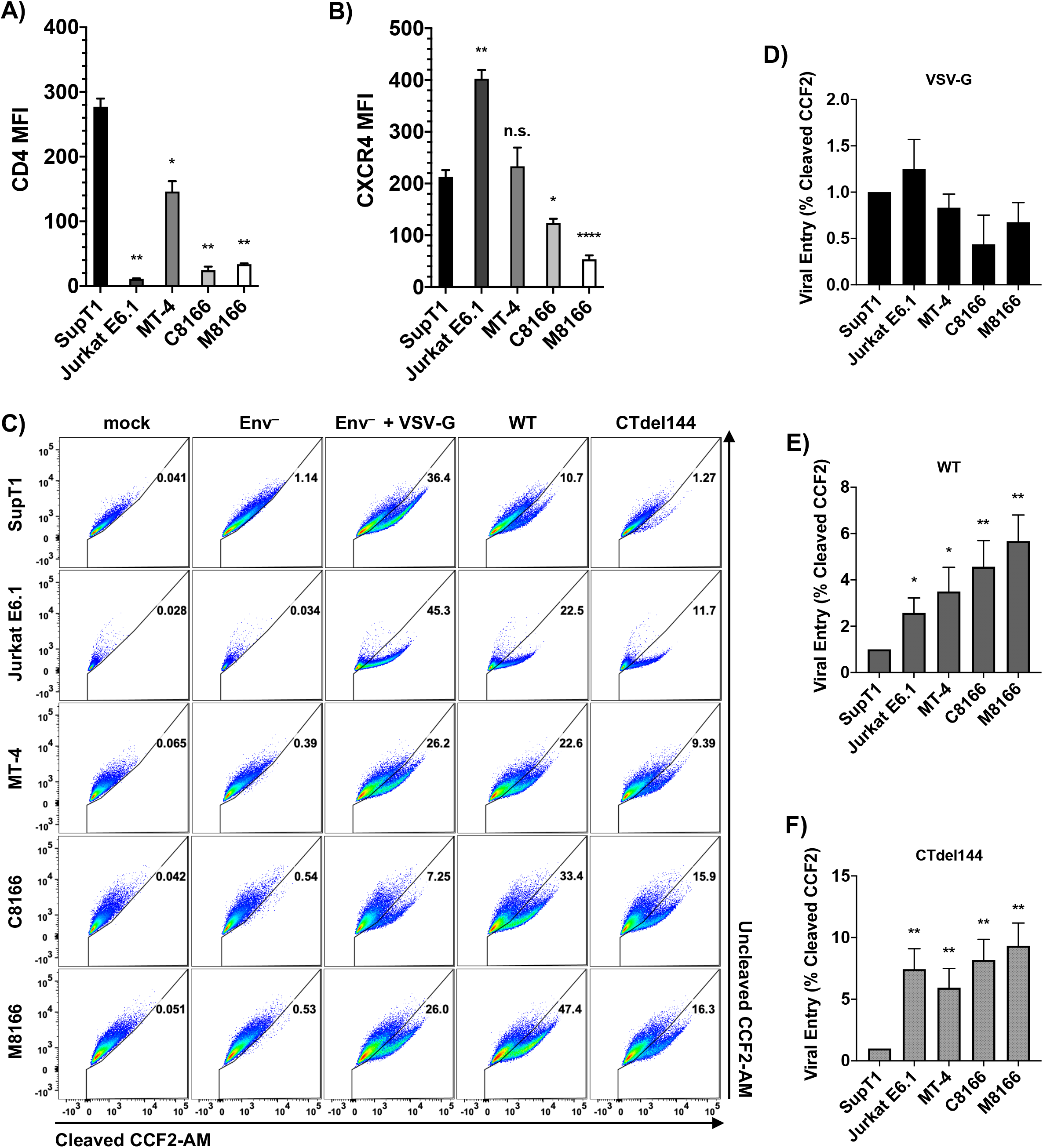
Viral entry mediated by full-length and CT-truncated Env is equivalent among HTLV-transformed T-cell lines. A) Surface CD4 and B) CXCR4 was measured by flow cytometry. MFI was determined by measuring the MFI value of CD4^+^ or CXCR4^+^ histograms and subtracting the isotype control MFI to directly compare MFIs between samples. To account for fluctuations in flow cytometry data that occur when comparing data from different experiments, cells from different acquisition dates were stained with antibody, fixed, and analyzed in parallel by flow cytometry. The bar graphs show the mean MFI values of cell surface A) CD4 and B) CXCR4, ± SD from three independent experiments and the bar shading for ease of comparison between panels A) and B). C) BlaM-Vpr-based viral entry assays were performed using NL4-3 expressing either WT or CTdel144 Env. A mock treatment and Env^-^ NL4-3 pseudotyped with VSV-G were used as a positive control and Env^-^ was used as a negative control. Representative FACS dot plots are shown. Virus dose used for each condition was titrated to avoid saturation (in the case of VSV-G and WT) and to be above the Env^-^ conditions (in the case of CTdel144). Therefore, values can only be compared between cell lines for each condition, not between conditions. The bar graphs show the fold change in viral entry relative to SupT1 (set at 1) between D) VSV-G-pseudotyped Env^-^ NL4-3, E) WT NL4-3, and F) CTdel144 NL4-3 with SD representing comparison of values derived from three independent experiments. Statistical significance was assessed by one-way ANOVA and Tukey’s multiple comparison test. P-values are defined in the Materials and Methods.

To determine whether increased efficiency of viral entry contributes to MT-4 permissivity, the β-lactamase (BlaM)-based viral entry assay was performed using virus bearing either VSV-G, WT HIV-1 Env, or the CTdel144 Env truncation mutant (Fig. 5C-F). Differences in viral entry mediated by VSV-G were not statistically significant between the cell lines (Fig. 5D). Entry mediated by WT HIV-1 Env was significantly lower in SupT1 than in the other T-cell lines, with MT-4 cells intermediate between Jurkat E6.1 and the other HTLV-1-transformed T-cell lines. Entry mediated by WT Env was not statistically different between MT-4 and Jurkat E6.1, C8166, and M8166 (Fig. 5E). Entry mediated by CTdel144 was equivalent among all cell lines tested, with the exception of SupT1, which displayed restricted entry consistent with low susceptibility to cell-free HIV-1 infection (6). These results demonstrate that the permissivity of MT-4 cells to the gp41 CT truncation mutant is not explained at the level of cell-free virus entry.

### Kinetics of virus release in MT-4 are faster than in other T-cell lines tested

Because the kinetics of HIV-1 replication in MT-4 cells are more rapid than in the other T-cell lines tested, we hypothesized that the kinetics of HIV-1 gene expression might also be faster in MT-4 cells. An equal number of cells were transduced with equivalent amounts of VSV-G-pseudotyped Env^−^ NL4-3 encoding eGFP in place of Nef (Fig. 6A-C). In this reporter virus, eGFP expression is under transcriptional control of the viral LTR; eGFP thus serves as a surrogate for LTR-mediated gene expression. Cells were analyzed for eGFP expression by flow cytometry at multiple time points post-transduction. We observed that a higher percentage of MT-4 cells expressed eGFP 24 hours post-transduction relative to the other T-cell lines (Fig. 6A). The percentage of MT-4 expressing eGFP increased ~3-fold 48 hours post-transduction, whereas the other cell lines tested exhibited a ~2-fold increase in the number of eGFP^+^ cells, with the exception of M8166, which exhibited only a minor increase in eGFP expression between 24 and 48 hours post-transduction. Therefore, the kinetics of HIV-1 protein production in MT-4 are faster relative to the other T-cell lines examined here.

**Figure 6.**
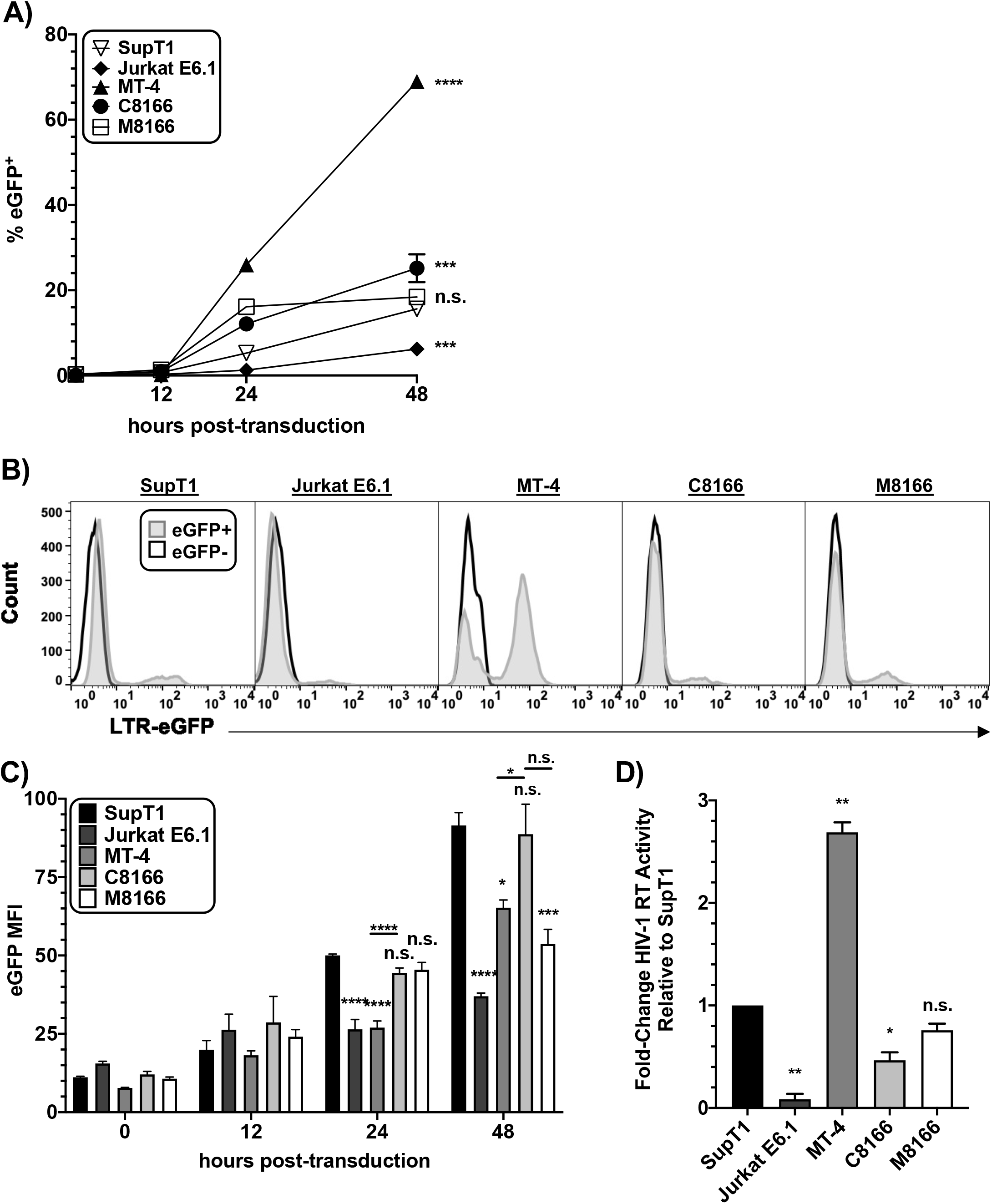
Kinetics of virus release in MT-4 are faster than in other T-cell lines tested. Cells were transduced with VSV-G-pseudotyped pBR43IeGFP-nef^-^Env^-^ reporter vector and collected at various time points. A) The percent of cells expressing eGFP at each time point was plotted. Error bars indicating the standard deviation of duplicate infections are too small to be seen. B) Histogram of eGFP expression in the cell lines 48-hour post-transduction indicating the percent of cells expressing eGFP and their MFI relative to an eGFP^-^ control. C) eGFP levels at various time points post-transduction (p.t.) were determined by the eGFP MFI from which the background MFI from the eGFP^-^ construct was subtracted. The bar graph shows the mean eGFP MFI, ± SD from three independent experiments. D) Cells lines were transduced with VSV-G-pseudotyped NL4-3 Env^-^, washed, and supernatant HIV-1 RT activity was measured 42-hours post-transduction. The bar graph shows the mean fold-change relative to the SupT1 reference line, ± SD from three independent experiments. Statistical analysis was performed to compare cell lines relative to SupT1 at individual time points post-transduction or between two samples as indicated by the horizontal line between two bars. n.s. indicates no statistical difference between the indicated cell line and the SupT1 reference or between two samples as indicated by the horizontal line between two bars. Statistical significance was assessed by one-way ANOVA and Tukey’s multiple comparison test.

Enhanced virus production and release on a per-cell basis could contribute to the gp41 CT-truncation permissive phenotype of MT-4 cells. To determine whether MT-4 express more HIV proteins than non-permissive cells, the eGFP median fluorescence intensity (MFI) was measured at different time points (Fig. 6B-C). At 24 hours post-transduction, the MFI of MT-4 was less than that of SupT1, C8166, or M8166. At 48 hours post-transduction, the MT-4 MFI increased to that of C8166 and M8166, indicating that the HIV gene expression on a per-cell basis between MT-4 and C8166 is equivalent, consistent with the data of Emerson et al (14). The finding that SupT1, a Tax^-^ non-permissive cell line, expressed more eGFP than MT-4 is inconsistent with the hypothesis that MT-4 are permissive to CT-truncation due to overall enhanced protein production per cell relative to non-permissive cells.

To explore the role of viral protein production kinetics in virus output, virus release in the T-cell panel was measured. Cells were transduced with Env^-^ VSV-G-pseudotyped NL4-3 and HIV-1 RT released into the supernatant was measured 42 hours post-transduction (Fig. 6D). Consistent with increased kinetics of HIV-1 protein expression compared to the other T-cell lines in the panel, HIV-1 RT production was highest for MT-4 compared to other cell lines. Taken together, these data indicate that on a per-cell basis, individual MT-4 cells do not exhibit higher steady-state cell-associated viral gene expression than the non-permissive cell lines tested here. However, MT-4 cultures release nearly three-fold more HIV-1 RT in a 42-hour period, indicating that the kinetics of HIV protein production in MT-4 cells are faster than in the other cell lines tested.

### Env incorporation in MT-4 is inefficient

Because Env incorporation into virus particles is essential for viral infectivity, we next investigated the role of the gp41 CT in Env incorporation in SupT1 and MT-4. Env incorporation was determined by two methods, radio-immunoprecipitation (radio-IP) and western blot (Fig. 7A-C). Equivalent numbers of both MT-4 and SupT1 cells were transduced overnight with VSV-G-pseudotyped HIV-1 expressing either WT or CTdel144 Env and washed extensively the following morning to remove unabsorbed virus. Cell- and virus-containing supernatants were collected, lysed, and prepared for analysis as described in the Methods. Consistent with our previous report (21), truncation of the gp41 CT in a non-permissive T-cell line, SupT1, resulted in an approximately 10-fold decrease in Env incorporation, as measured by gp120 and gp41 levels in virions, compared to WT (Fig. 7A). In contrast, in MT-4 cells, truncation of the gp41 CT resulted in an approximately 2-fold reduction in Env incorporation (Fig. 7B). We next tested whether higher levels of WT Env are incorporated into virions produced in MT-4 compared to SupT1 cells. Virus lysates were similarly prepared as in Fig. 7A-B, RT-normalized, subjected to SDS-PAGE, and Env band intensities were measured (Fig. 7C). Surprisingly, MT-4 were found to incorporate less WT Env than SupT1. To gain more insight into why MT-4 incorporate less Env than the non-permissive SupT1 cell line, various parameters of viral assembly were determined (Fig. 7D-F). Env processing in MT-4 was lower than in SupT1 (Fig. 7D). The ratio of cell-associated gp120 to total Gag was also lower for MT-4 than SupT1 (Fig. 7E). Consistent with a previous report that examined other non-permissive T-cell lines (14), we observed that Gag processing in MT-4 is more efficient than in the non-permissive SupT1 (Fig. 7F). More rapid Gag processing is likely an indication of more rapid Gag trafficking, membrane association, and/or assembly, perhaps resulting in assembly of the Gag lattice before Env is recruited (19).

**Figure 7.**
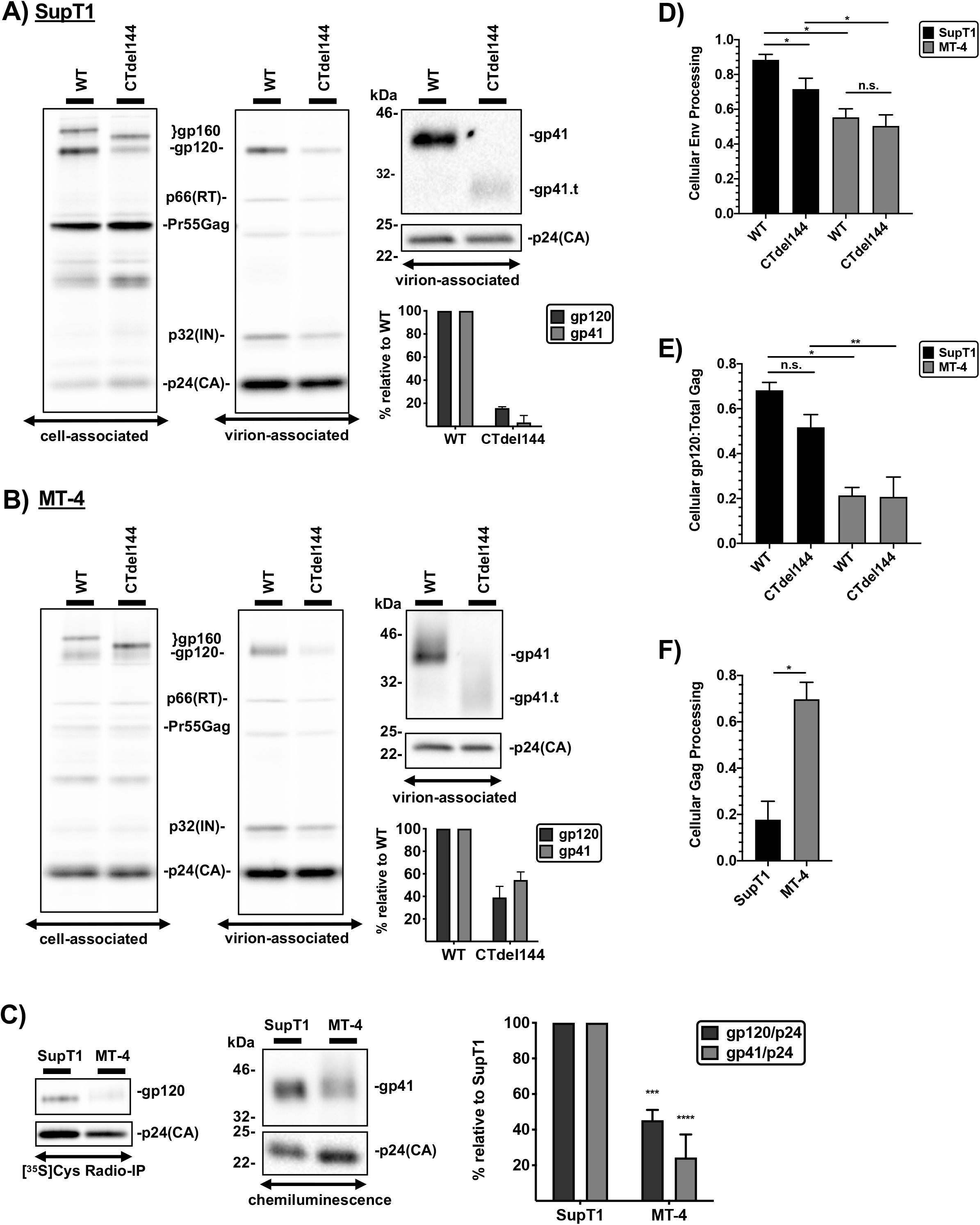
Env incorporation in MT-4 cells is inefficient. SupT1 (A) and MT-4 (B) cells were transduced with RT-normalized VSV-G-pseudotyped NL4-3 encoding either WT or CTdel144 Env. Cells were metabolically labeled with [^35^S]Cys and cell and virus lysates were immunoprecipitated to detect HlV proteins. The locations of p66(RT), the Gag precursor Pr55Gag, p32(IN), and p24(CA) are indicated. Western blotting was performed on the virus fraction to detect gp41 and p24(CA) using equal amounts of WT and CTdel144 viral lysates. gp41.t indicates the position of the truncated gp41, CTdel144. The fold-change in Env incorporation between WT and CTdel144 Env, calculated by determining the ratio of virion-associated gp41 to p24(CA) relative to the WT condition, is indicated below the western blots. C) SupT1 and MT-4 were transduced as in (A-B) and RT-normalized virus was used to compare gp41 content in the virus fraction. Samples were analyzed by [^35^S]Cys radio-immunoprecipitation (IP) and western blot (detected by chemiluminescence). The bar graph shows the mean values relative to the SupT1 reference line, ± SD from three independent experiments. D) Env processing efficiency was quantified by dividing cell-associated gp120 by the total cell-associated Env band intensity [gp120/(gp120+gp160)]. E) The cell-associated Env-to-Gag ratio was determined as [gp120/(Pr55Gag+p24)]. F) Gag processing was determined by dividing p24(CA) by total Gag band intensity. D-F) The bar graphs show the mean values, ± SD from three independent experiments. n.s. indicates no statistical difference between two samples as indicated by the horizontal line. C-E) Statistical significance was assessed by one-way ANOVA and Tukey’s multiple comparison test or F) paired Student’s t-test.

### High infectivity of virions produced from MT-4 does not explain their permissivity to *gp41* CT truncation

The surprising result that Env incorporation in MT-4 is inefficient relative to SupT1 suggested that there is something inherently more efficient about viral transmission in the MT-4 cell line. To determine whether cell-free virions produced from MT-4 cells are more infectious than virions produced from the other cell lines in our panel, TZM-bl infectivity assays were performed (Fig. 8A-C). Cells were transduced with VSV-G-pseudotyped HIV-1 encoding either WT or CTdel144 Env and collected 42 hours post-transduction. RT-normalized virus supernatants from the T-cell panel were used to infect TZM-bl cells in parallel. Consistent with our findings that Env incorporation of the CT-truncated Env is reduced ~10-fold and ~2-fold in SupT1 and MT-4, respectively, cell-free infectivity was also reduced to a similar extent by the CTdel144 mutation (Fig. 8A). Thus, under these conditions and with these cell lines, cell-free infectivity closely correlates with levels of virion-associated Env.

**Figure 8.**
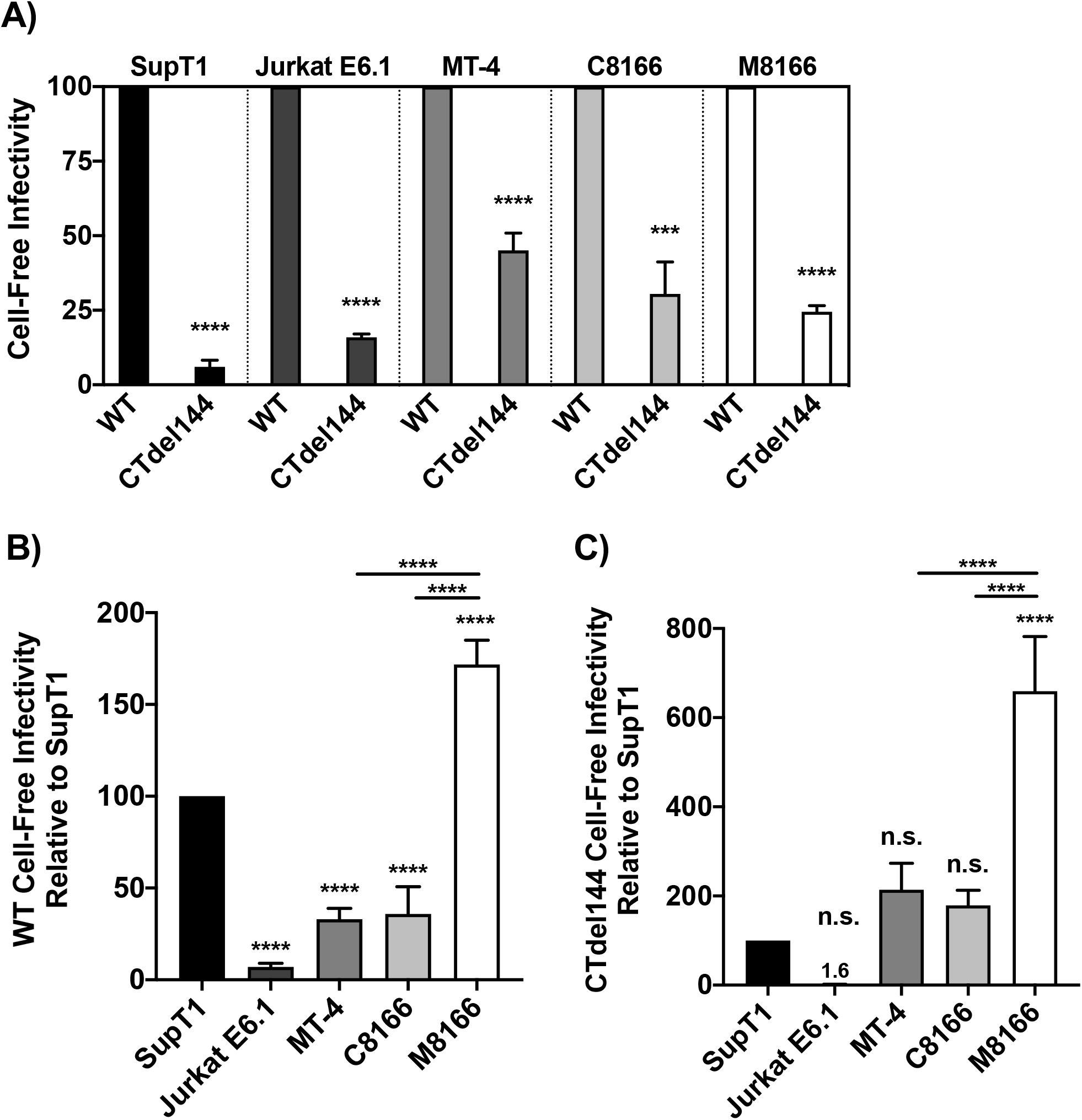
High infectivity of virions produced from MT-4 cells does not explain their permissivity to gp41 CT-truncation. Cells were transduced with VSV-G-pseudotyped NL4-3 encoding either WT or CTdel144 Env. Viral supernatant was collected 42 hr post-transduction. Supernatants were RT normalized, and a serial dilution of virus was used to infect TZM-bl cells. Luciferase values were then used to determine the relative infectivity of virus produced from each T-cell line. A) The bar graph shows the mean values of CTdel144 relative to WT (set at 100), ± SD from three independent experiments. B-C) The same virus was used as in A) to compare WT and CTdel144 virus infectivity between cell lines. The bar graphs show the mean values of B) WT and C) CTdel144 relative to the SupT1 reference line (set at 100), ± SD from three independent experiments. Shading of individual columns indicates values for the individual cell lines for ease of comparison between data sets in A-C. Statistics relative to the SupT1 reference line, or between two samples as indicated by the horizontal line between two bars, are shown. “n.s.” indicates no statistical difference between the indicated cell line to the SupT1 reference. Statistical significance was assessed by A) Student’s t-test and B) one-way ANOVA and Tukey’s multiple comparison test.

To directly compare the infectivity between WT and CTdel144 virus produced from all five T-cell lines, TZM-bl cells were infected in parallel with the same RT-normalized virus produced as in Fig. 7A. SupT1 were used as the standard of comparison. We observed the infectivity of WT virus produced by all cells, with the exception of M8166, to be significantly lower than WT virus produced by SupT1 (Fig. 8B). Interestingly, CTdel144 virions produced in the non-permissive cell line M8166 exhibited ~3-fold higher levels of infectivity compared to MT-4 and C8166 (Fig. 8C), suggesting that cell-free infectivity is, at most, a minor contributing factor to the CT-permissive phenotype of MT-4.

### MT-4 express higher levels of cell surface CT-truncated Env than other T-cell lines tested

Because Env incorporation in MT-4 is reduced compared to the other T-cell lines tested, we next investigated whether Env is efficiently expressed on the cell surface of MT-4. To measure cell surface-associated Env, the T-cell panel was transduced with VSV-G pseudotyped WT or CTdel144 NL4-3 and fixed 42 hours later, a time point at which the cells would be expected to express HIV-1 proteins, but not display extensive syncytium formation. Fixed cells were then fluorescently labeled with anti-gp120 antibodies and Env expression was measured by flow cytometry as a shift in the histogram relative to an Env^−^ control (Fig. 9A). WT Env expression on MT-4 cells was higher than on the other cell lines tested (Fig. 9B). Consistent with the role of the gp41 CT in regulating endocytosis (73–76), truncation of the gp41 CT enhanced cell-surface Env expression in all cell lines including MT-4, although in M8166 the increase was not statistically significant (Fig. 9B).

**Figure 9.**
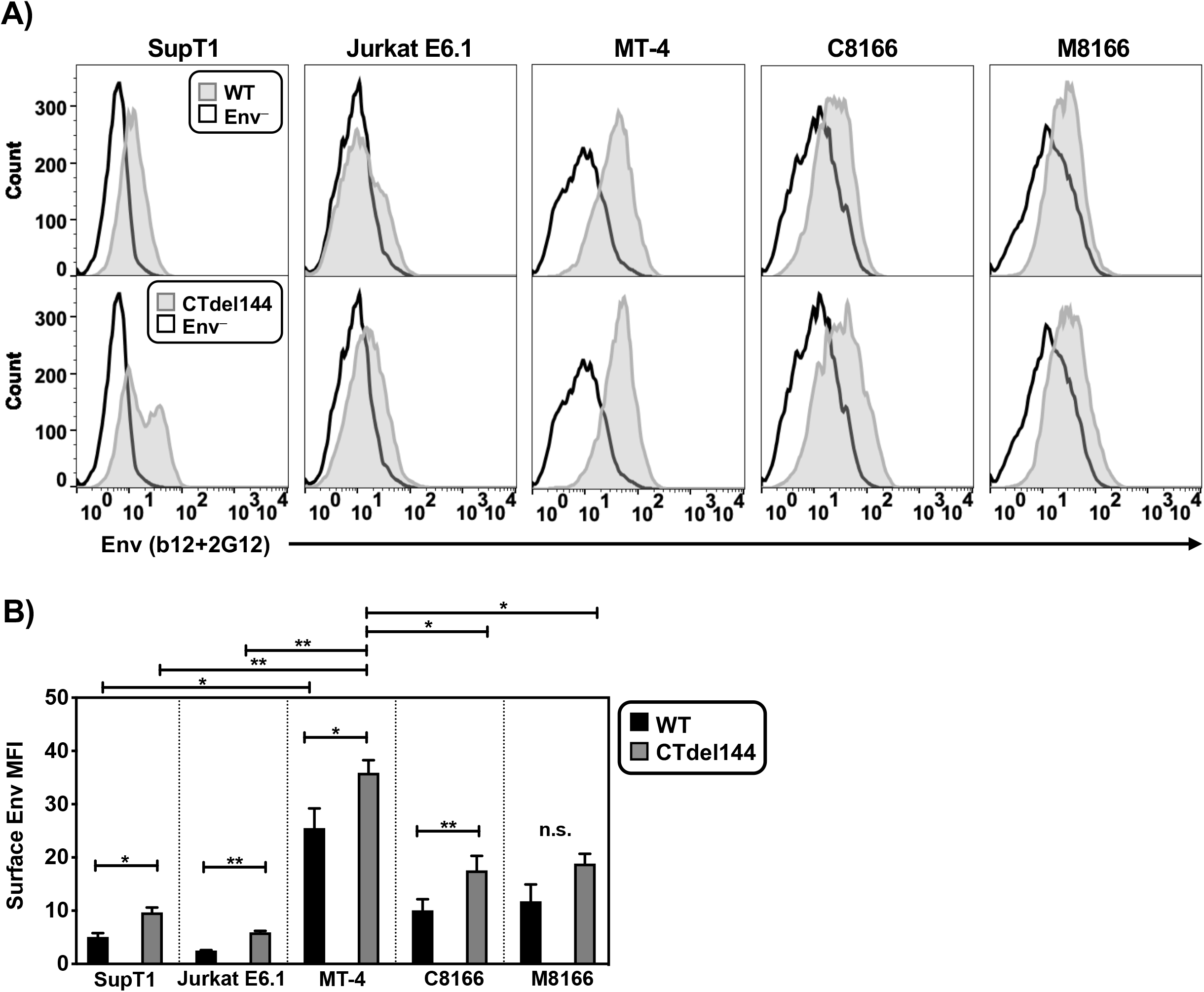
MT-4 cells express higher levels of surface CT-truncated Env than other T-cell lines tested as determined by flow cytometry. VSV-G-pseudotyped NL4-3 encoding either WT or Ctdel144 Env, or no Env (Env^-^) was used to transduce T cells. 40 hours post-transduction, cells were fixed and stained with anti-Env antibodies for analysis by flow cytometry. A) Histogram of surface Env for each cell line is shown. Env^-^ cells were used as a control for background staining. B) Surface Env MFI was determined by measuring the MFI value of the Env^+^ histogram and subtracting the Env^-^ MFI to directly compare MFIs between samples. To account for fluctuations in flow cytometry data that occur when comparing data from three independent experiments, cells were stained with antibody and analyzed in parallel by flow cytometry. The bar graph shows the mean values of WT and CTdel144 Env MFI, ± SD from three independent experiments. n.s. indicates no statistical difference between WT and CTdel144 for the M8166 cell line. Error bars ± SD from 3 independent experiments. Statistical significance was assessed by Student’s t-test.

The enhanced surface expression of Env observed in MT-4 cells could be explained by either defects in Env internalization or high levels of viral protein production relative to the other cell lines, resulting in high levels of cell-associated Env. Having established that MT-4 do not exhibit higher levels of LTR-mediated gene expression than some other cell lines in the panel, we sought to determine whether Env internalization is defective in MT-4. To confirm the higher levels of cell-associated Env on MT-4 relative to the nonpermissive SupT1 cell line, MT-4 and SupT1 cells were transduced with VSV-G-pseudotyped HIV-1, pulse-chased with a fluorescently labeled anti-gp120 monovalent Fab probe and analyzed by confocal microscopy (Fig. 10A) (19). A monovalent Fab fragment was used to probe for Env levels to avoid bivalent antibody crosslinking of Env trimers, which has been shown to induce Env internalization (77). The anti-gp120 probe used in the pulse-chase assay labeled both the internalized pool of Env and surface Env. The total (non-biosynthetic) Env was quantified by combining both the internalized pool and surface Env values. As a control to establish whether AP-2-dependent internalization machinery was functional, the cell media were supplemented during the short anti-gp120 pulse with fluorescent transferrin (Tfn) to label the endosomal compartments (78, 79) (Fig. 10A – uninfected conditions). Tfn was internalized in both SupT1 and MT-4, confirming that both cell lines have an intact endocytosis machinery.

**Figure 10.**
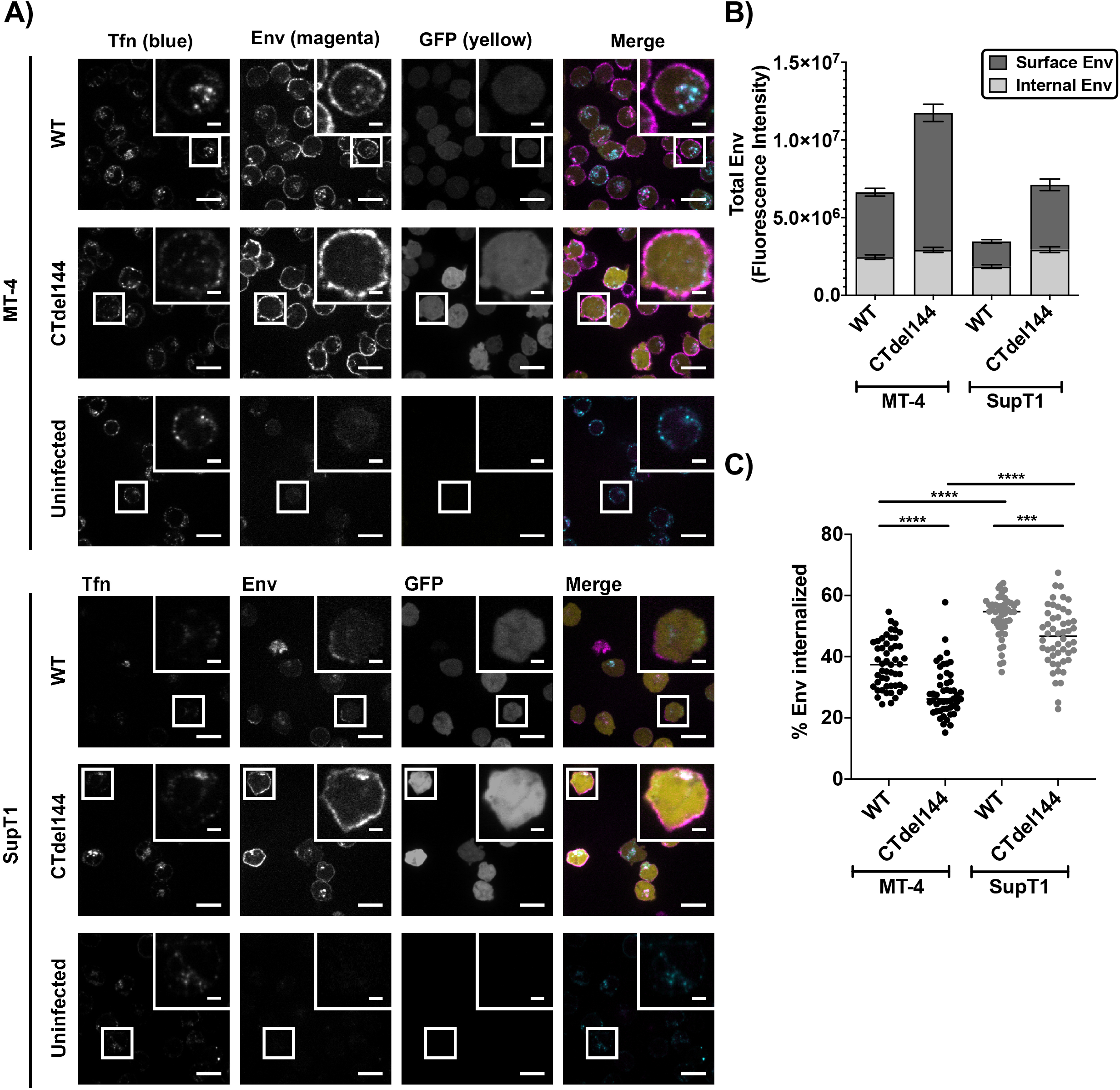
MT-4 cells express higher levels of Env than SupT1, which is further enhanced by truncating the gp41 CT, as determined by confocal microscopy. MT-4 or SupT1 cells were infected with a replication-and-release-defective HIV-1 mutant (described in methods) expressing an eGFP reporter (yellow), and either WT or CTdel144 Env. Approximately 40 hours post-infection, cells were simultaneously pulsed with anti-Env Fab b12-Atto565 (magenta) and Transferrin-AF647 (blue) for 12 min and then chased for 50 min. Cells were fixed and imaged by confocal fluorescence microscopy. A) Representative confocal slices of transduced MT-4 or Jurkat E6.1 cells after the pulse-chase labeling. Scale bars 15 μm, inset scale bars 3 μm. B) Total surface Env and total internal Env per cell were measured using the integrated fluorescence intensity for regions defining the PM of the cell (surface) and a region defining the interior of the cell (internal), using a maximum intensity projection through a 4.5 μm confocal range centered in the approximate middle of a single cell. n = 50 infected cells per sample. The bar graph shows the mean values of Surface and Internal Env levels. C) Percent Env internalization was calculated as the percent of internal Env above the total Env signal (internal and surface). The scatter plot shows the mean values of percent internalized Env. n = 50 infected cells per sample. Error bars ± SEM. Statistical significance was assessed by one-way ANOVA and Tukey’s multiple comparison test.

To determine whether differences exist in Env internalization in SupT1 and MT-4, the ratio of surface to internalized Env was analyzed (Fig. 10B-C). Consistent with the flow cytometric analysis (Fig. 9), MT-4 expressed higher levels of total and surface Env than SupT1 (Fig. 10B). Surface Env expression on MT-4 was further increased by truncating the gp41 CT. To gain further insight into the ability of MT-4 to internalize Env, we determined the percent Env internalization during the pulse-chase (Fig. 10C). WT Env was internalized more rapidly in SupT1 than in MT-4. As expected, truncation of the gp41 CT resulted in less internalized Env for both MT-4 and SupT1. Altogether, the data suggest that Env levels on the surface of MT-4 cells may be higher than on other lines due to slower internalization.

### MT-4 serve as better targets for C-C transmission than the other cell lines tested

The results presented above suggest that levels of virion-associated Env and cell-free particle infectivity do not account for the permissivity of MT-4 cells to CT-truncated Env. We therefore explored the role of C-C transmission in the gp41 CT-truncation permissive phenotype. To gain a better understanding of MT-4 cells as a target for HIV-1 infection, we compared the relative ability of cell lines in the T-cell panel to be infected via C-C transmission, using 293T cells as the donor. Non-lymphoid 293T cells do not express the adhesion molecules LFA-1, ICAM-1, or ICAM-3 (80). Using 293T cells as the virus-producing donor cell enabled the study of viral transmission independent of differences in LFA-1:ICAM1/3 engagement, surface Env levels, and kinetics of HIV release, as C-C transfer in this system is dependent primarily on Env-CD4 interactions. Because ~50% of our Jurkat E6.1 cells do not express CD4 (Fig. 5A), this cell line was excluded from the analysis as the CD4^-^ cells would not become infected in the assay. The vector used in this analysis encodes eGFP under control of the HIV-1 LTR; eGFP therefore labels cells infected by either the cell-free or C-C route. 293T were transiently transfected with an HIV-1 proviral clone encoding eGFP and 24 hours later either cocultured with dye-labelled T cells or overlaid with a transwell containing dye-labelled T cells. Viral transfer in the co-culture thus represents the summation of both cell-free and C-C infection events (Fig. 11A). Evaluation of the transwell data showed that cell-free infection of MT-4 is less efficient relative to C8166 and M8166, but more efficient than SupT1 (Fig. 11B). Subtracting the transwell from the co-culture values produced the C-C contribution (Fig. 11C). The results indicated that MT-4 cells were more efficiently infected by C-C transmission than SupT1 or C8166. High MT-4 and M8166 susceptibility as target cells is not due to higher CD4 expression, since we found that SupT1 express higher levels of CD4 compared to either MT-4 or M8166 (Fig. 5A) and equivalent or greater levels of CXCR4 than MT-4 or M8166, respectively (Fig. 5B). The differences in susceptibility to C-C transmission from 293T donor to MT-4 and M8166 targets were not statistically significant, suggesting that high susceptibility to C-C transmission likely contributes to, but does not entirely account for, the CT-truncation permissive phenotype of MT-4.

**Figure 11.**
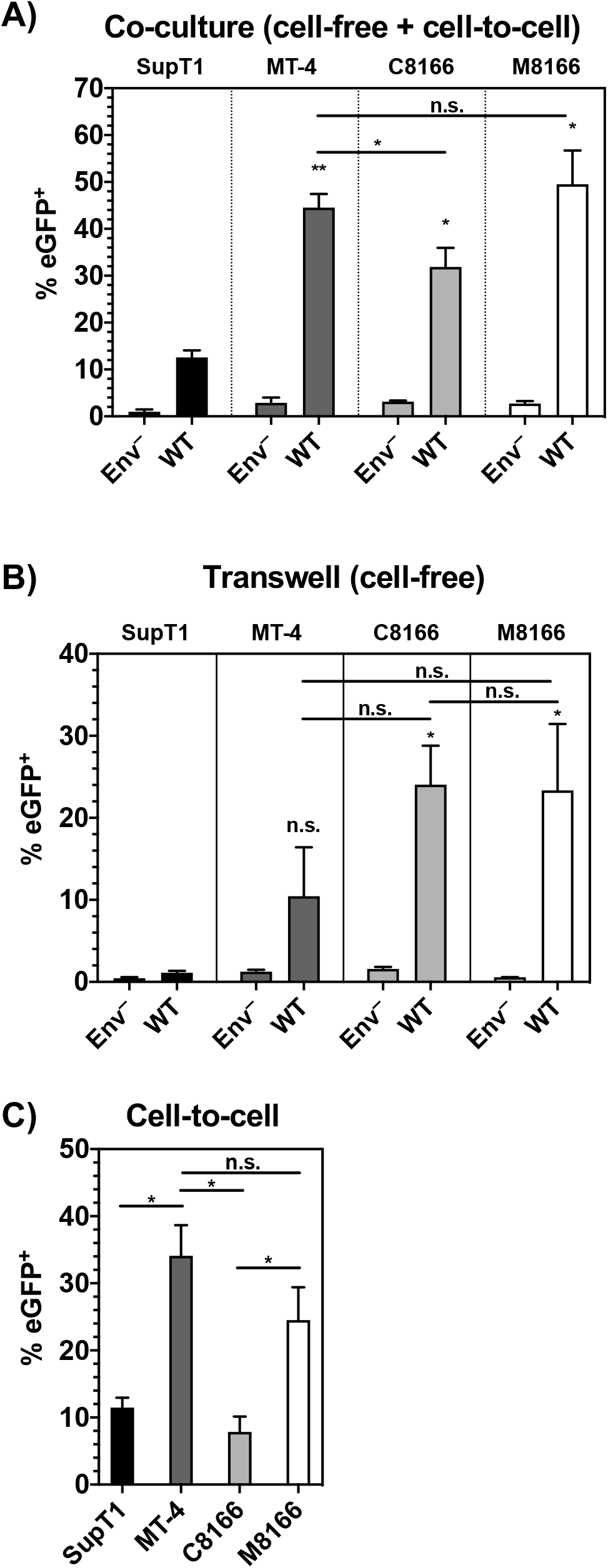
MT-4 serve as better targets for C-C transmission than the other cell lines tested. 293T cells were transfected with pBR43IeGFP-nef^-^ encoding either WT Env or Env^-^. Twenty-four hours post-transfection, dye-labelled T cells were either co-cultured or added to a transwell exposed to the 293T supernatant. Eighteen hours post-co-culture, BMS-806 was added to prevent multiple cycles of infection and the formation of syncytia. Forty-four hours post initial co-culture, cells were collected, fixed, and analyzed by flow cytometry. A-B) The percentage of cells expressing eGFP was determined and plotted. C) The transwell value was subtracted from the co-culture value to determine the contribution of cell-to-cell transmission. The bar graphs show the mean values of the percent of cells positive for eGFP expression for the A) co-culture, B) transwell (cell-free), and C) C-C transmission, ± SD from three independent experiments. Shading of individual columns indicates values for the cell line for ease of comparison between data sets. n.s. indicates no statistically significant difference between the indicated cell line and the SupT1 reference or between two samples as indicated by the horizontal line connecting two bars. Statistical significance was assessed by one-way ANOVA and Tukey’s multiple comparison test.

## Discussion

It has been previously reported in a number of studies that the CTdel144 gp41 truncation mutant is unable to establish a spreading infection in most T-cell lines or primary hPBMCs. In this study, we sought to elucidate the basis for the unusual, and in our analysis unique, permissivity of the MT-4 T-cell line to truncation of the gp41 CT. To this end, a panel of validated T-cell lines known or reported to be permissive or non-permissive to gp41 CT truncation was compared to MT-4. We confirmed that the MT-4 T-cell line is permissive to CT-truncation, but observed that other HTLV-transformed T-cell lines previously reported as permissive (14, 58, 81) are not. Differences in methodology or cell line origin likely explain the contrast in these findings. In part because of these differences, we extensively validated the T-cell lines used in the study. We found that HTLV-I Tax expression, viral entry efficiency, cellular levels of viral protein expression, virion-associated Env content, and cell-free viral infectivity did not solely account for permissivity to gp41 CT truncation. A combination of rapid HIV-1 gene expression, enhanced cell-surface Env expression, and high susceptibility to C-C transmission compared to the non-permissive lines tested in this study likely explains the ability of MT-4 to overcome the requirement for the gp41 CT.

It was previously suggested that MT-4 express higher levels of Gag proteins after HIV-1 infection relative to the non-permissive cells examined (14). We found that in MT-4 cultures, more cells were eGFP^+^ over time indicating faster HIV-1 protein production kinetics compared to non-permissive cells. HTLV-I Tax expression and upregulated NF-κB activity were not associated with greater HIV-1 protein production; C8166 and M8166 expressed comparable levels of eGFP per cell relative to Tax^−^ SupT1 and Jurkat E6.1. These contrasting results are likely explained by methodological differences; our flow cytometry approach allowed us to quantify both eGFP expression per cell, and number of cells expressing eGFP, at various time points, while the western blot approach used by Emerson et al. (14) measured total HIV-1 Gag expression in the cell cultures at a single time point. Both our study and that of Emerson et al. found that MT-4 cells release approximately 3-fold more virus than non-permissive cells. We conclude that this is due to faster kinetics of HIV-1 protein production in MT-4 cells rather than more protein being expressed per infected cell, relative to non-permissive cells.

A recent report found that MT-4 and C8166 are likely permissive to type I IN mutations due to HTLV-I Tax expression inducing NF-κB protein recruitment to the HIV-1 LTR on unintegrated HIV-1 DNA (46). If this phenomenon were the cause of T-cell line permissivity to truncation of the gp41 CT, Tax^+^ C8166 and M8166 would also be permissive to gp41 CT truncation. Furthermore, we found that MT-4, C8166, and M8166 all display higher NF-κB activity compared to SupT1, indicating that they are all in a hyper-NF-κB activated state. Therefore, while Tax expression was found to overcome type I IN defects and activate transcription of non-integrated HIV-1 DNA in C8166 and MT-4 (46), this phenomenon does not account for the unique ability of MT-4 to host multiple rounds of CTdel144 infection.

Reduced Env incorporation in MT-4 was associated with a commensurate reduction in cell-free infectivity, suggesting that viral particles produced by MT-4 are inherently less infectious than those produced by non-permissive cells. We also observed that MT-4 cells exhibit a higher level of Gag processing relative to SupT1 and a lower level of Env processing. More efficient Gag processing may be a consequence of increased assembly kinetics. Previous work has led to the hypothesis that the timing of Gag assembly versus Env expression on the surface can influence the efficiency of Env incorporation; a delay in Env trafficking to the surface relative to completion of Gag assembly reduces the efficiency of Env trapping by the nascent Gag lattice (19), a phenomenon that could account for the relatively inefficient Env incorporation observed in MT-4 relative to SupT1.

Consistent with the role of the gp41 CT in regulating Env internalization from the PM, truncation of the gp41 CT resulted in enhanced surface Env expression on all T-cell lines tested (with the exception of M8166 in which the increase was not statistically significant) as observed in previous studies (17, 82, 83). Analysis of Env internalization, using a pulse-chase assay, found that the rate of Env internalization was reduced in MT-4 compared to SupT1. This reduced Env internalization may contribute to the reduced Env incorporation seen in MT-4, consistent with a role for gp41 CT-dependent internalization from the PM in Env incorporation during viral assembly (15–17). This finding suggests a recycling-independent Env incorporation model in MT-4, wherein the presence of a full-length tail allows for some enhancement of WT Env incorporation by Gag lattice trapping during particle assembly (22), compared to the passive, less efficient incorporation of the gp41 CT-truncated mutant which is unable to be trapped. The higher surface expression of Env in MT-4 cells contributes to the smaller effect of CT truncation in Env incorporation in MT-4 relative to non-permissive T-cell lines. This is consistent with the observation that in HeLa cells the Env incorporation defect observed with CTdel144 could be overcome by increasing Env expression (15); more Env on the surface leads to an increase in non-specific (passive) incorporation that is less dependent on the gp41 CT and its trapping by the Gag lattice (1). The higher surface expression of Env in MT-4 relative to non-permissive T-cell lines may also contribute to the formation of VSs and more productive transfer of CTdel144, which is more fusogenic than Env with a full-length CT (18), by C-C transmission.

Our approach of using 293T as virus-donor cells allowed us to test the susceptibility of the T-cell line panel to infection by either cell-free or C-C transmission independent of the ability of the T-cell lines to serve as donor cells. We observed an approximately 3-fold higher susceptibility of MT-4 and M8166 cells to C-C transmission from 293T donors compared to SupT1 and C8166. These data indicate that a differential susceptibility to C-C transmission between C8166 and M8166 (a subclone of C8166 more susceptible to formation of syncytia) arose during the generation of M8166. Because the lipid composition of the target cell membrane has been shown to affect viral fusion and entry (84, 85), it is possible that differences in PM lipid composition between the cell lines studied here account for the high susceptibility of MT-4 and M8166 to C-C transmission. Further studies will be required to evaluate this possibility.

This study reveals that multiple factors are associated with the ability of the MT-4 T-cell line to support the replication of a gp41 CT truncation mutant. The requirement for the gp41 CT in MT-4 is likely overcome by the additive effects of rapid HIV-1 protein production, high levels of cell-surface Env expression, and increased susceptibility to CC transmission compared to non-permissive cells. These results highlight that Env incorporation, viral transfer, and ultimately the establishment of a spreading infection are influenced by a number of factors including the kinetics of viral protein expression and virus assembly, the levels of Env expressed on the cell surface, and the rate of Env trafficking and internalization.

## Materials & Methods

### Cell lines and culture

293T [obtained from American Type Culture Collection (ATCC)] and TZM-bl [obtained from J. C. Kappes, X. Wu, and Tranzyme, Inc. through the NIH AIDS Reagent Program (ARP), Germantown, MD] cell lines were maintained in DMEM containing 5% or 10% (vol/vol) fetal bovine serum (FBS), 2 mM glutamine, 100 U/mL penicillin, and 100 μg/mL streptomycin (Gibco) at 37 °C with 5% CO_2_. Jurkat E6.1, MT-4, C8166-45 (referred to as C8166), and M8166 were obtained from Arthur Weiss (Cat# 177), Douglas Richman (Cat# 120), Robert Gallo (Cat# 404), and Paul Clapham (Cat# 11395), respectively, through the NIH ARP. The source of SupT1 used in this study is unknown. The SupT1-CCR5 (86) T-cell line was a generous gift from James Hoxie. The ATL Tax^−^ cells lines (ED, ATL-35T, and TL-Om1) were a generous gift from Chou-Zen Giam. T-cell lines were maintained in RPMI-1640 medium containing 10% FBS, 2 mM glutamine, 100 U/mL penicillin, and 100 μg/mL streptomycin (Gibco) at 37 °C with 5% CO_2_. Whole blood was obtained from healthy donors via the NIH Clinical Center. hPBMCs were isolated using a ficoll gradient and stimulated with 2 μg/mL PHA-P for 3 – 5 days before infection, then cultured in 50 U/mL IL-2.

### Cloning and plasmids

The HIV-1 pNL43-nef^−^-eGFP reporter vector (also called pBR43IeGFP-nef^−^; Cat# 11351) [obtained through the NIH ARP from Frank Kirchhoff]. The Env^−^ construct was generated by restriction digest of the pNL4-3/KFS clone (87), referred to here as pNL4-3 Env^−^, and target vector with StuI and XhoI restriction enzymes followed by ligation of the Env fragment into pBR43IeGFP-nef^−^ to generate pBR43IeGFP-nef^−^Env^−^. Unless otherwise indicated, the full-length HIV-1 clade B molecular clone pNL4-3 was used (88). The Env^−^ (87) and CTdel144 clone (26) were described previously. Wild-type and CTdel SIVmac239 were generated by Bruce Crise and Yuan Li and were a generous gift from Jeff Lifson. Plasmid sequences were confirmed by restriction digest with HindIII and Sanger sequencing. The HIV-1 YU2 Vpr β-lactamase expression vector (pMM310) was [obtained from Michael Miller, Cat # 11444, through the NIH ARP]. The plasmids pHR’CMV-GFP and pHR’CMV-Tax were a generous gift from Chou-Zen Giam.

For confocal microscopy, non-propagating, release-defective constructs were generated (pSVNL4-3-ctfl-dPol-dVV-GFP-3’LTR and pSVNL4-3-ctfl-dPol-dVV-dCT-GFP-3’LTR where ctfl = C-terminal FLAG tag on Gag, dPol = deletion of Pol, dVV = deletion of Vif and Vpr, by removal of the Aflll fragment, GFP = green fluorescent protein in place of Nef). Expression plasmids for Env and associated mutants in addition to Gag were natively expressed from the reference HIV-1 clone NL4-3 with the following modifications: the pNL4-3 vector was sub-cloned into an SV40 ori-containing backbone (pN1 vector; Clontech; pSVNL4-3), deletion of *pol* by removal of the BclI-NsiI fragment, mutation of the p6 PTAP motif (455PTAP458-LIRL) (89), addition of a coding C-terminal FLAG tag to the Gag open reading frame to detect Gag expression (GSDPSSQ500-SGDYKDDDDK) (90) and removal of the 5’ portion of the *nef* open reading frame and replacement with a GFP coding sequence.

### Preparation of virus stocks

293T cells were transfected with HIV-1 proviral DNA using Lipofectamine 2000 (Invitrogen) according to the manufacturer’s instructions. Virus-containing supernatants were filtered through a 0.45-μm membrane 48 h post-transfection and virus was quantified by measuring RT activity. VSV-G–pseudotyped virus stocks were generated from 293T cells co-transfected with proviral DNA and the VSV-G expression vector pHCMV-G (91) at a DNA ratio of 10:1.

For confocal microcopy, virus particles were made using the pSVNL4-3 plasmids. Briefly, VSV-G-pseudotyped, single-round viruses were produced by transfecting 293T cells with the pSVNL4-3 plasmids, the psPAX2 packaging plasmid (a gift from Didier Trono, Addgene plasmid #12260), and pVSV-G expression plasmid using PEI transfection reagent (Alfa Aesar/Thermo Fisher Scientific). The virus was harvested at ~48 hours after transfection and stored at −80 °C.

### STR profiling and cell line validation

The identity of cells in the T-cell line panel was confirmed by performing STR profiling as described previously (43). Briefly, genomic DNA was extracted and sent to Genetica (LabCorp) for profiling. The URL for this data set is: https://amp.pharm.mssm.edu/Harmonizome/gene_set/IRF3/ENCODE+Transcription+Factor+Targets. The obtained STR profile was compared to the Cellosaurus reference STR using the % Match formula (92).

### Illumina RNA-Seq

RNA was extracted from T cells in their exponential growth phase using QIAshredder (Qiagen, Cat# 79654) and RNeasy Plus Mini Kit (Qiagen, Cat# 74134). RNA sample integrity was assessed by determining the RNA Integrity Number (RIN) with an Agilent 2100 Bioanalyzer instrument and applying the Eukaryote Total RNA Nano assay. RIN values were between 9 and 10, indicating intact RNA. Samples were sequenced on a Hiseq 2500, generating an average of 50 million raw reads per sample. Transcript sequence reads were normalized against the total reads for each cell line to generate a reads per kilobase per million mapped reads (RPKM) value. The RPKM is a relative measure of transcript abundance.

A panel of NF-κB-dependent genes was obtained by random selection of genes from a database of NF-κB target genes (70). An IRF3-dependent gene panel was obtained from the Harmonizome search engine (71) by searching the ENCODE Transcription Factors gene set titled “IRF3” (72). This gene set contains 4159 IRF3 transcription factor target genes obtained from DNA-binding by ChIP-seq datasets. Genes dually targeted by NF-κB and IRF3 were removed from the IRF3-dependent gene panel used in our analysis.

To analyze expression of NF-κB and IRF3 target genes, the average RPKM for each parent gene transcript was mined from the total acquired RNA-seq data set. RPKM values of zero were set to 1 to allow for fold-change analysis. HTLV-transformed cell lines were compared to the lymphoma-derived SupT1 line to generate the fold-change value. This value was then converted by Log2 transformation.

### Virus Replication Assays

Virus replication kinetics were determined in Jurkat E6.1 and MT-4 cell lines as previously described (26). Briefly, T cells were transfected with proviral clones (1 μg DNA/10^6^ cells) in the presence of 700μg/ml DEAE-dextran. MT-4, C8166, and M8166 cells were split 1:2 every 2 days, and SupT1 and Jurkat E6.1 were split 1:3 every 2-3 days with fresh media. VSV-G-pseudotyped virus was used to inoculate stimulated hPBMCs. Virus stocks were normalized by ^32^P RT activity and used to initiate spreading infection. After a 2-hour incubation with VSV-G-pseudotyped virus, cells were washed and resuspended in fresh RPMI-10% FBS. Every other day half the medium was replaced without disturbing the cells. Virus replication was monitored by measuring the RT activity in collected supernatants over time. RT activity values were plotted using GraphPad Prism to generate replication curves.

### HIV-1 infection of T cells

293T cells were plated and co-transfected via lipofectamine the next day with the pNL4-3 proviral clone and the VSV-G expression vector, pHCMV-VSV-G, at a 10:1 ratio. 48 hours post-transfection, supernatants were passed through a 0.45 μm filter and RT activity was measured by ^32^P RT activity assay as described (93). T cells were plated the night before infection in fresh media at a density of 5×10^6^ cells / 2 mL. The following day, cells were infected overnight with RT-normalized virus.

### Western blotting for viral proteins

The morning after transduction, cells were washed extensively to remove any unabsorbed virus, and plated in 1.5 mL of RPMI-10. 42 hours post-infection, cells were pelleted and lysed. Virus-containing supernatants were passed through a 0.45μm filter. 10 μL were set aside for RT assay and 200 μL were set aside for TZM-bl infectivity assay. The remaining filtered virus-containing supernatant was layered on a 20% w/v sucrose/phosphate buffered saline (PBS) solution and spun for 1.25 h at 41,500 x *g* at 4°C in a Sorvall S55-A2 fixed angle rotor (ThermoFisher Scientific). Cell and virus fractions were lysed in lysis buffer (30 mM NaCl, 50 mM Tris-HCL pH 7.5, 0.5 % Triton X-100, 10mM Iodoacetamide, complete protease inhibitor (Roche)). Lysates boiled with 6x loading buffer (7 mL 0.5 M Tris-HCL/0.4 % SDS, 3.8 g glycerol, 1 g SDS, 0.93 g DTT, 1.2 mg bromophenol blue) were subjected to SDS-PAGE on 12% 1.5 mm gels and processed using standard western blotting techniques. All antibodies were diluted in 10 mL of 5% milk in TBS blocking buffer. HIV proteins were detected with 10 μg/mL polyclonal HIV immunoglobulin (HIV-Ig) obtained from the NIH ARP. Anti-human IgG conjugated to horseradish peroxidase (HRP) was obtained from SigmaAldrich (Cat# GENA933) and used at a 1:5000 dilution. gp41 was detected with 2 μg/mL10E8 monoclonal antibody obtained from the NIH ARP (Cat# 12294) followed by anti-human IgG-HRP as above.

To detect HTLV-I Tax protein, 2×10^6^ cells were lysed in 100 μL of lysis buffer and 30 μL of 6x loading buffer and boiled for 10 min. 20 μL of cell lysates were loaded on a 12% 1.5mm Tris-glycine gel and processed using standard western blotting techniques. HTLV-I Tax was detected with an anti-Tax-1 mouse monoclonal (Abcam, Cat# ab26997) at a 2 μg/mL concentration followed by goat-anti-mouse-HRP antibody (Thermo Fisher, Cat # 32230) at a 0.5 μg/mL concentration. β-actin was used as a loading control and detected using anti-β-actin conjugated directly to HRP (Abcam, Cat# ab49900). Protein bands were visualized using chemiluminescence with a Bio-Rad Universal Hood II ChemiDoc and then analyzed with ImageLab v5.1 software or the Azure Spot Analysis Software.

Metabolic labeling and radioimmunoprecipitation were performed as previously described (21). Phosphor screens were imaged with a Bio-Rad Personal Molecular Imager System and quantitative analysis of bands was performed with the Azure Spot Analysis Software.

### Viral entry assay

BlaM-Vpr-containing viruses were produced by transient co-transfection of 239T cells with pNL4-3 constructs encoding either Env^−^, WT, or CTdel144 sequences, a second plasmid (pMM310 – NIH ARP Cat# 11444) encoding the BlaM-Vpr fusion protein, and a third plasmid (pAdvantage - Promega, Cat# E1711) to enhance transient protein expression at a 6:2:1 ratio. For the VSV-G control, pHCMV-VSV-G was provided in *trans* at a 10:1 ratio relative to pNL4-3. Virus-containing supernatant was filtered 48 hours post-transfection and concentrated using Lenti-X concentrator (TakaraBio, Cat# 631231) according to the manufacturer’s instructions. After concentration, the virus particles were resuspended in CO_2_-independent media (ThermoFisher, Cat# 18045088), aliquoted, and stored at −80°C. Resuspended virus was diluted 10,000x and virus concentration was measured using two methods: (1) p24^gag^ concentration by Lenti-X GoStix Plus (TakaraBio, Cat# 631280) and (2) ^32^P RT activity (93).

Entry of BlaM-Vpr-containing virus was detected using a fluorescent substrate, CCF2-AM (Thermo Fisher, Cat# K1032) (94). To perform the BlaM-Vpr assay, a suspension of 20×10^6^ cells/mL was made. 100 μL of this suspension was aliquoted per well of a U-bottom plate. Cells were infected for 4 h at 37°C with 400 ng p24^gag^. After incubation with virus, cells were washed 2x with CO_2_-independent media supplemented with 10% FBS. Cells were then resuspended in 100 μL CO_2_ independent media with 10% FBS and 20 μL of the CCF2-AM reagent (prepared according to the manufacturer’s instructions). Cells were incubated for 1 hour with the CCF2-AM reagent in darkness at room temperature. Cells were then washed 2x with PBS, resuspended in CO_2_-indpendent media + 10 % FBS and left at room temperature in darkness overnight. Sixteen hours later, cells were fixed in 100 μL 4% PFA and analyzed the same day. Flow cytometric analysis utilized a Fortessa X-20 flow cytometer (BD Bioscience) and FlowJo software (Tree Star Inc, Ashland, OR, USA).

### Single-cycle infectivity assays

TZM-bl infectivity assays were performed as previously described (95). Briefly, 20,000 TZM-bl cells were plated in a flat-bottomed white-walled plate (Sigma Aldrich, Cat# CLS3903-100EA). The following day, the cells were infected with serial dilutions of RT-normalized virus stocks in the presence of 10 μg/mL DEAE-dextran. Approximately 36 hours post-infection, cells were lysed with BriteLite luciferase reagent (Perkin-Elmer) and luciferase was measured in a Wallac BetaMax plate reader.

### Flow cytometry

Cells were resuspended in 8% BSA/PBS at a concentration of 10^7^ cells/mL. 100 μL was aliquoted into a 96-well V-bottom plate (SigmaAldrich, Cat# CLS3897). An antibody solution was made using 20 μL of either isotype or target antibody per test, as recommended by the manufacturer. The following antibodies from BD Pharmigen were used: APC mouse IgG2a **κ** isotype control (Ca# 555576), APC mouse anti-CD184 (CXCR4) (Cat# 560936), PE mouse anti-CD4 (Cat # 555347). The isotype control for CD4 was acquired from Biolegend (PE mouse IgG1, **κ** isotype control, Cat# 400113). Cells were incubated with antibodies for 20 min at room temperature, then washed 3x with PBS. Samples were analyzed via the BD FACSCalibur flow cytometer (BD Bioscience) and FlowJo software (Tree Star Inc.).

To detect cell-surface Env, infected cells were washed with PBS and fixed with 4% PFA overnight at 4°C. The following day, the PFA was quenched with 0.3 M glycine in PBS using three rounds of 5 min incubations. Cells were then resuspended in 100 μL of 8% BSA/PBS. An antibody cocktail was made by diluting b12 and 2G12 in 8% BSA/PBS [obtained from Dr. Dennis Burton and Carlos Barbas, and Polymun Scientific, respectively, via the NIH-ARP] and aliquoting 100 μL to each sample to obtain a final concentration of 2 μg/mL of both antibodies. The cell:antibody mixture was incubated at room temperature for 20 min. Unbound antibody was removed with three PBS washes. Goat anti-human conjugated to Alexa Fluor 488 was used as a secondary antibody (Invitrogen, Cat# A11013) at a concentration of 2 μg/mL. Cells were incubated with secondary antibody for 20 min at room temperature and washed 3x with PBS before analysis on the BD FACSCalibur.

### C-C transmission assay

293T were transfected with pBR43IeGFP-nef^+^ (Cat# 11349; NIH ARP) or an Env^-^ derivative (described above). 24 hours post-transfection, dye-labelled T cells were added to the transfected 293T cells at a 4:1 ratio in either a co-culture or to the top layer of a transwell. 12 hours post-coculture, BMS-806 was supplemented into the media at a concentration of 2000 nM to prevent the formation of syncytia and T cell to T cell transmission. Forty-eight hours post-coculture, cells were fixed in 4% PFA and analyzed by flow cytometry. Data were collected via CellQuest and processed with FlowJo software (Tree Star Inc).

### Production of anti-Env monovalent Fab b12-Atto565 probe

The anti-Env b12 Fab fragment recombinant expression vector was a kind gift from Dennis Burton (96). A 4x lysine tag, to enhance dye conjugation efficiency (97), was added to the C-terminus of the b12 light chain using site-directed mutagenesis. Expression of b12 was carried out as previously described (19, 98). Briefly, clarified cell lysates were purified by protein G affinity chromatography (Gold Biotechnology, Inc.). The b12 Fab, typically 99% pure, was conjugated with Atto565 N-hydroxysuccinimidyl ester (Sigma-Aldrich). Typically, a labeling ratio of 1 Atto565 dye molecule per b12 Fab molecule was achieved.

### Pulse-chase staining

For each sample, 6-8×10^5^ MT-4 or SupT1 cells were transduced with VSV-G-pseudotyped virus encoding either WT or CTdel144 Env (described above). Approximately 40 hours post-infection, the cells were blocked with 10% BSA in complete media for 30 min at 37°C with 5% CO_2_, stained with custom Fab b12-Atto565 (25 nM) and Transferrin-AlexaFluor647 (Transferrin-AF647; Invitrogen; 25 μg/mL) in media with 6% BSA, and washed thrice for 5 min with the same media. The cells were then placed onto 18 mm coverslips pre-treated with poly-L-lysine (Ted Pella Inc.), fixed with 4% and 0.2% paraformaldehyde and gluteraldehyde in PBS, respectively. Coverslips were then mounted onto glass slides with Fluoromount-G (Southern Biotech).

### Microscopy

Imaging of the cellular specimens was performed with a spinning-disk confocal microscope built on an inverted Nikon Ti-E base (Solamere Technology Group Inc., Salt Lake City, UT) using a 60x, CFI Plan Apo Lambda 1.4 NA oil-immersion objective (Nikon Instruments). Fiber-coupled lasers (OBIS, Coherent) were used with a CSU-X A1 spinning disk unit (Yokogawa Electronics) to excite and collect confocal fluorescence sections. Transferrin-AF647 was imaged using a 640 nm laser at 12 mW with 40 ms exposure, Atto565 was imaged using a 561 nm laser at 24 mW with 100 ms exposure, and GFP was imaged using a 488 nm laser at 12 mW with 40 ms exposure. Laser power was measured at the specimen. Z-stacks for each field of view were collected at 0.3 μm spacing.

### Image analysis

Quantitation of surface-exposed and internalized Env signal in the pulse-chase assay images was performed using ImageJ software. First, for each z-stack, a maximum intensity projection was generated for a 4.5 μm range through the middle of the cells. The background was subtracted using ImageJ’s built-in “rolling ball” background subtraction process with a radius of 150 pixels. Cells were excluded from analysis if they did not express GFP (non-infected cells) or did not internalize transferrin (dead or dysfunctional cells), or if the maximum intensity projection did not represent an equatorial section of the whole cell. Integrated intensity was then estimated using two regions of interest (ROIs) in the Atto565 channel: (1) an outer ROI was manually segmented to enclose the entire cell, including any plasma-membrane-associated signal, and (2) an interior ROI was manually segmented just inside the PM, excluding any PM-associated signal. Signal for Env on the PM was thus enclosed between these two ROIs, and signal for the internalized Env pool was enclosed within the interior ROI. The integrated intensity of the outer ROI (1) minus that of the inner ROI (2) served as a measure of surface-exposed Env, while the integrated intensity of the inner ROI (2) served as a measure of internalized Env.

### Statistics

Statistics were calculated using GraphPad Prism version 8 for Mac OS (GraphPad Software, La Jolla, CA). Unpaired Student’s t-tests were performed and two-tailed *P < 0.05, **P < 0.01, ***P < 0.001, and ****P < 0.0001 were considered statistically significant. GraphPad Prism was also used to calculate standard error and to assess statistical significance by one-way ANOVA. P values for Student’s t-test and one-way ANOVA analysis are defined with the same cut-offs.

### Ethics Statement

PBMCs were obtained from anonymous, de-identified blood donors to the NIH Department of Transfusion Medicine Blood Products Program (NIH CC-DTM).

## List of abbreviations

CT: cytoplasmic tail
VS: virological synapse
C-C: cell-to-cell
Env: envelope glycoprotein
STR: short tandem repeat
MFI: median fluorescence intensity
PM: plasma membrane
Radio-IP: radio-immunoprecipitation
hPBMC: human peripheral blood mononuclear cells
RT: reverse transcriptase
VSV-G: Vesicular stomatitis virus glycoprotein
PBS: phosphate buffered saline
FBS: fetal bovine serum
WT: wild type
HRP: horseradish peroxidase
ER: endoplasmic reticulum
MA: HIV-1 Matrix protein
EBV: Epstein-Barr virus
HTLV: Human T-cell leukemia virus
ALL: Acute lymphocytic leukemia
ATL: Adult T-cell lymphoma
ATCC: American Type Culture Collection
NIH ARP: NIH AIDS Reagent Program
n.s.: not statistically significant
IN: HIV-1 Integrase protein
CA: HIV-1 Capsid protein
RIN: RNA Integrity Number

## Declarations

### Authors’ contributions

MVF and EOF designed the study. MVF and HKH performed the experiments and analysis, and NP generated the fluorescently labelled b12 monovalent Fab for the microscopic analysis. PT performed initial western blots comparing Env content in virions produced in SupT1 and MT-4. SBvE contributed valuable feedback and guidance on the microscopy experiments. All authors made contributions to writing the manuscript.

## Acknowledgements

We thank members of the Freed and van Engelenberg labs for helpful discussion and for critical review of the manuscript. We thank James Hoxie for providing the SupT1-CCR5 cell line, Erin M. Hall at Genetica (Lab Corp) for assistance with STR profiling and data interpretation, Douglas Richman for providing historical information on the treatment of MT-4 cells prior to donation to the NIH ARP, and James A. Thomas for assisting with the BlaM-Vpr flow cytometry acquisition. We thank Chou-Zen Giam for helpful discussion and for providing the ATL Tax^-^ T-cells (ED, ATL-55T, and TL-Om1) and expression vectors for Tax and GFP expression. We also thank Yongmei Zhao at the NCI-Center for Cancer Research Sequencing Facility for performing the Illumina RNA-seq and assisting with the differential RNA analysis. This study was supported by the Intramural Research Program of the Center for Cancer Research, National Cancer Institute, NIH and by an Intramural AIDS Research Fellowship (for MVF).

